# IGF-like growth factors that couple food sensing to developmental decisions employ different release mechanisms in a single pair of *C. elegans* chemosensory neurons

**DOI:** 10.64898/2026.01.23.701173

**Authors:** Julia M Perhacs, Lauren Klabonski, Mingjie Ying, Yair Argon, Tali Gidalevitz

## Abstract

Sensory neurons modulate organismal physiology and behavior in part by releasing neuropeptides and neurotrophins or growth factors, *via* the dense core vesicle (DCV) pathway. The precise matching of the sensory input to the identity of released vesicle cargo, and the timing and location of its release, is necessary to ensure appropriate responses. In some neurons, several neuropeptides or growth factors can be co-produced simultaneously, but found in distinct populations of vesicles. While this may permit their release independently of each other, in response to different stimuli, the responsible molecular pathways are not well understood. In *C. elegans*, the chemosensory ASI neuron pair expresses multiple neuropeptides and growth factors, including the structurally homologous *C. elegans* IGF-like growth factors DAF-28, INS-6, and INS-4, whose release in young larvae couples food sensing to the commitment to reproductive development. We find that these growth factors require separate molecular pathways for their release. While the axonal release of INS-6 protein required the Calcium-dependent Activator Protein for Secretion (CAPS/UNC-31), the release of DAF-28 was CAPS-independent. Consequently, the function of endogenous *daf-28*, but not that of *ins-6* and *ins-4*, is independent of *unc-31*. This difference is unexpected, as CAPS/UNC-31 is thought to control the regulated pre-synaptic/axonal release of DCV cargoes in *C. elegans* neurons, and demonstrates a divergence in vesicle release mechanisms. In addition, we find that mechanisms for delivering vesicles to the axons are also divergent: while neuropeptide NLP-21 was dependent on the clathrin adaptor AP-3 for its selective localization to axons, four insulin/IGF-like growth factors tested - DAF-28, INS-6, INS-4, and INS-22, were AP-3-independent for either axonal localization or function. Our data uncover molecular divergence in the pathways controlling axonal localization and release of neuropeptides and growth factors, including in a single neuron. These divergent vesicle pathways provide new means for the immediate and tunable changes in neuronal outputs, in addition to the known, less immediate mechanism of differential transcriptional regulation of DCV cargoes. Future delineation of the molecular composition of these pathways is necessary to understand how neurons match organismal responses to specific sensory inputs.

**Graphical Abstract:** Molecular divergence in the pathways controlling axonal localization and release of neuropeptides and growth factors

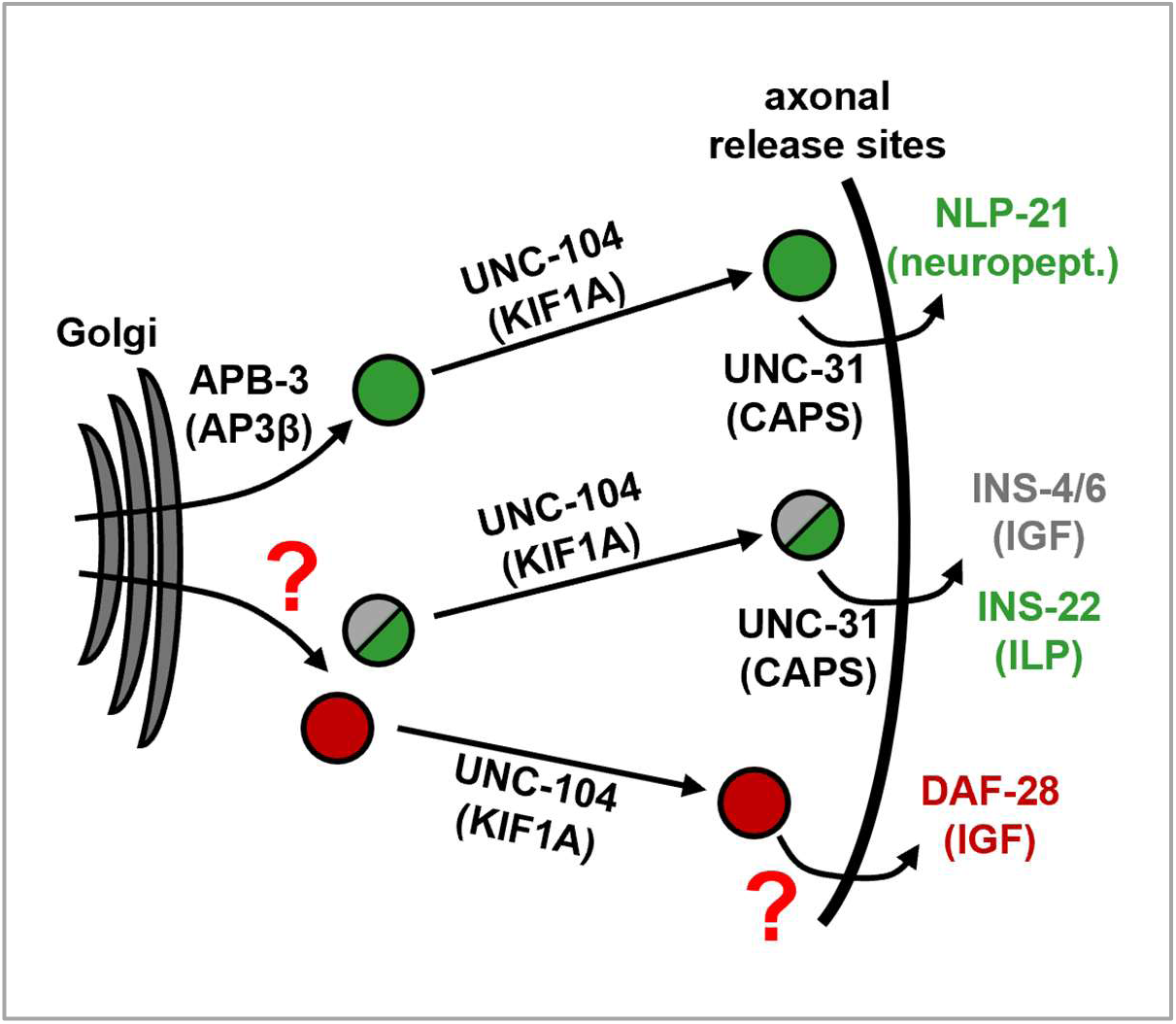

## Introduction

To ensure organismal survival and adaptation, sensory neuronal systems must detect and correctly respond to variable environments. In addition to electrical or chemical synaptic transmission, sensory neurons transmit information by releasing neuromodulators, neuropeptides, and neurotrophins or growth factors, *via* the dense core vesicle (DCV) pathway. The peptide and protein DCV cargoes are made in the ER and Golgi, processed and transported to their release sites through the regulated pathway^1–6^, and released by vesicle fusion either at pre-synaptic terminals, peri-synaptically, or at extrasynaptic sites^7–11^. They then act locally or at a distance^12,13^, to modulate neurotransmitter signaling and to control various organismal functions and behaviors^11,14^. Defects in the DCV pathway are associated with developmental, neurological, and neuropsychiatric disorders^15–22^.

Certain neurons can simultaneously express several neuropeptides and/or growth factors, sometimes with divergent or even antagonistic functions^23–26^. Some of these co-produced signaling molecules share vesicles^27,28^, while others, including examples of neuropeptides cleaved from a single precursor protein, are targeted into distinct, non-overlapping vesicle populations^29–35^. The targeting of different cargos to non-overlapping vesicle populations provides a potential for independent regulation of their release^35,36^. This may be crucial in animals with smaller nervous systems, such as *C. elegans*, where a small number of neurons needs to discriminate and appropriately respond to multitude of environmental inputs. Indeed, several examples of independent release of co-produced neuropeptides have been reported in *C. elegans*, including in the ASI sensory neuron, in response to different monoamines, and in AVK and AIY interneurons, in response to an autocrine loop or oxidative stress, respectively^24,26^. Such differential release of DCV cargoes, along with their differential transcription, serves to increase the repertoire and selectivity of neuronal responses, and impart flexibility to anatomically hard-wired circuits^36,37^. However, mechanisms generating the divergence in vesicle pathways are only beginning to be mapped.

One example for differential release mechanisms involves modulating Ca^2+^ sensitivity of different vesicle populations *via* recruitment of different isoforms of synaptotagmin (Syt), required for the Ca^2+^-dependent membrane fusion^38–42^. Because Ca^2+^ signals are local and often have different properties, they may selectively activate fusion proteins possessing different affinities for Ca^2+43,44^. In the mouse olfactory bulb neurons, the growth factor IGF-1 and the neuropeptide ANF are found in two distinct, non-overlapping populations of vesicles^32^; Syt10, but not co-expressed Syt1, was found to localize only to the IGF-1 – containing vesicles and to selectively control its release and the resulting olfactory behaviors^32,45,46^. It is unknown how soluble cargos like IGF-1 or ANF, which are confined to the vesicle lumen, can be paired to the specific synaptotagmin isoforms, or how other cytosolic proteins could differentially regulate the release of soluble cargoes.

Furthermore, to be differentially regulated by local Ca^2+^, the neuropeptides or growth factors would need to be targeted to distinct release locations. The differentially released neuropeptides in *C. elegans* ASI neurons are released from different sites in the axon and cell body^24^; this is also true for the differential release of *Drosophila* insulin-like peptide 2 in clock neurons^47^, and of BDNF from axons *vs* dendrites in mammalian neurons^39,48^. However, while several mechanisms for targeting soluble cargoes into the regulated pathway are known, including cargo receptors, and Ca^2+^- and pH-driven aggregation or phase-separation^49–52^, how these cargoes are then targeted to distinct release sites within the axon, or between axons, cell bodies, and dendrites, is unknown. This is in contrast to the much better understood mechanisms for differential targeting of transmembrane cargos. For example, membrane proteins destined for the axon are enriched into the corresponding vesicles through preferential interaction of their cytosolic domains with AP3 clathrin adaptor complexes, which then leads to their trafficking into the axon by appropriate kinesin motors^1,53–55^. Soluble neuropeptides and growth factors, in contrast, cannot directly access this cytosolic machinery.

*C. elegans* neurons have been used extensively to dissect DCV biology^1,2,4,56–59^, owing to the high conservation and low genetic redundancy of relevant pathways, and the accessibility of genetic and imaging approaches. Furthermore, the uniform microtubules orientation in both dendrites and axons^60^ simplifies the analysis of axonal *vs* dendritic polarization mechanisms. *C. elegans* exhibit sophisticated behavioral and developmental responses to changes in their environment, driven in part by the release of neuronal growth factors and neuropeptides^61–63^. A decision in the early larval stage to either commit to reproductive development or enter dauer diapause is one such response, controlled by the bilateral amphid chemosensory neurons, including the ASI^64^. Upon sensing pro-growth conditions - high ratio of food to population density (pheromone), and low temperature, the ASI neuron pair releases three insulin/IGF-like growth factors, DAF-28, INS-4, and INS-6; their release signals commitment to reproductive development and prevents dauer entry^65–69^. All three are agonists of DAF-2, the worm’s insulin-like growth factor 1 receptor (IGF1R) homologue, and belong to the same β-type family of insulin/IGF-like peptides (ILPs), based on their disulfide bonds arrangement^70^. DAF-28 is the predominant signal from the ASI under the continuous presence of pro-growth environment, but all three can partially compensate for each other, since their combined deletion causes unconditional dauer entry, and re-expression of either one reverses this defect; in addition, INS-6 and INS-4 are released from the ASJ and motor neurons upon re-appearance of food, to promote exit from dauer^67,68,71,72^. Besides these three IGFs, the ASI neurons at the time of dauer decision express several other modulators, including the TGFβ protein DAF-7, that promotes development and prevents dauer^73,74^, and the insulin homolog INS-1 with the opposite function - it promotes, rather than prevents dauer^67,75^. We have previously found that, unlike DAF-28, which localizes to the axons, DAF-7 is released from the sensory dendrite^76^. Thus, in addition to the transcriptional control of these growth factors^67,71,73,74,77,78^, the ASI neurons must differentially control their release, to couple the sensing of the environment to developmental plasticity.

We find that despite their structural and functional homology, the insulin/IGF-like INS-4, INS-6, and DAF-28 are released *via* two distinct mechanisms from *C. elegans* neurons. Our genetic data show that the function of endogenous *ins-4* and *ins-6* depends on the conserved DCV exocytosis priming factor, CAPS/UNC-31, as is expected for DCV cargo^57,79^. However, *daf-28* function is UNC-31-independent. In agreement, the functional fluorescent fusions of DAF-28 and INS-6 proteins are differentially dependent on UNC-31 for their release from the ASI axons. DAF-28 does utilize the regulated pathway, as both DAF-28 and INS-6 are actively and selectively localized to their release sites *via* axonal kinesin KIF1a/UNC-104. Because trans-membrane DCV proteins require the AP3 clathrin adaptor complex for their KIF1a-dependent axonal localization^53^, we asked if this is also true for these non-membrane soluble cargos. We found that neither the function of endogenous *daf-28*, *ins-4*, and *ins-6*, nor the polarized localization of DAF-28 and INS-6 proteins to the ASI axons depend on AP3β/APB-3. Similarly, polarized axonal localization of another insulin/IGF-like protein, INS-22, a known DCV cargo^4^, was AP3β-independent in motor neurons. However, in the same motor neurons, axonal targeting of neuropeptide NLP-21, also a known DCV cargo^4^, was strongly dependent on AP3β. Our data thus uncover distinct vesicle pathways in *C. elegans* neurons that diverge in both the initial targeting of the soluble cargo to the axon, and in the release mechanisms, and show that neurons have a striking ability to distinguish between structurally similar insulin/IGF-like DAF-28, INS-4, and INS-6.

## Results

### DAF-28/IGF does not require CAPS/UNC-31 for its function in promoting development, unlike structurally and functionally similar INS-4 and INS-6

Neuronal release of IGF-like proteins DAF-28, INS-4 and INS-6 in young larvae (late L1 (larval stage one) or early L2) signals commitment to reproductive development and prevents dauer entry. Previous work demonstrated that the three proteins are redundant for signaling developmental commitment, because *daf-28(-)*;*hpDf761* animals that lack all three IGFs entered dauer even in pro-growth conditions, while ectopic re-expression of either one of the three was sufficient to reverse the dauer entry^67,68^. For DAF-28 and INS-6, re-expression in the ASI and ASJ sensory neurons was sufficient for the rescue, while INS-4 seemed to act in the GABAergic motor neurons^68^.

We took advantage of their functional redundancy to ask whether DAF-28, INS-4 and INS-6 are released through the same canonical DCV mechanism. Mammalian CAPS proteins, CAPS1 and 2, are thought to control dense-core vesicle exocytosis in neurons and neuroendocrine PC12 cells^80–83^, and the sole *C. elegans* CAPS protein, UNC-31, was previously shown to be required for the neuronal release of model DCV cargoes^4,57,84,85^. Thus, we asked whether loss of UNC-31 (*e928* null allele^57,86^) in *daf-28(-)* animals, whose development depends on the function of INS-4 or INS-6, *vs.* in *hpDf761*animals, whose development depends on the function of DAF-28, causes inappropriate dauer entry, as would be expected if the remaining growth factors required UNC-31 (scheme in Fig. 1B). The *hpDf761* deficiency deletes *ins-4*, *ins-5* and *ins-6* genes, but since *ins-5* does not contribute to the dauer decision^68^, we will henceforth use *ins-4/6(-)* designation for clarity.

**Fig. 1.**
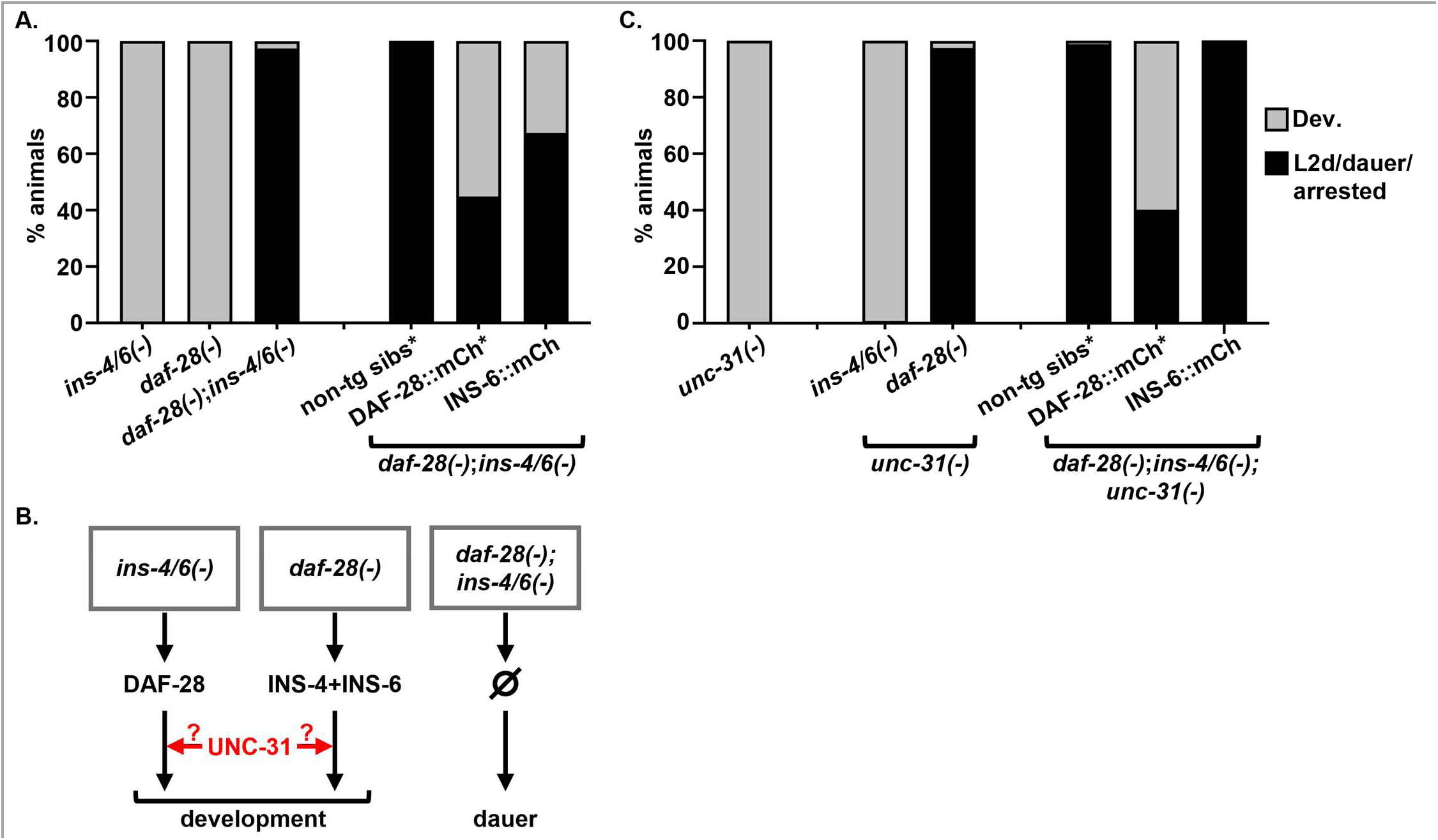
Endogenous DAF-28 does not require CAPS/UNC-31 for its function during dauer decision, unlike structurally and functionally similar INS-4 and INS-6. A. Percent animals entering reproductive development (gray) or in L2d/dauer/L1 arrest (black) at 46-48 hours post embryonic gastrula stage. L1 arrest was no more than 3.5% (see Data Suppl.). Star indicates DAF-28::mCherry and non-transgenic siblings growing on the same plates, scored separately. B. Schematic of the functional assay to determine UNC-31-dependency of endogenous DAF-28, INS-4, and INS-6. Mutant genotypes, proteins still expressed in the neurons, and the expected phenotypes are indicated. C. Endogenous DAF-28 and transgenic DAF-28::mCherry support development in the absence of UNC-31, while endogenous INS-4, INS-6, and INS-6::mCherry do not. Scored as in A, L1 arrest was no more than 3.5% (see Data Suppl.).

As reported^68^, both *ins-4/6(-)* and *daf-28(-)* animals developed normally, while almost all *daf-28(-)*;*ins-4/6(-)* animals entered dauer under pro-growth conditions of abundant food, low population density, and non-stressful growth temperature (20⁰C) (Fig. 1A). Loss of UNC-31 caused ∼98% abnormal dauer entry in *daf-28*-deficient animals (Fig. 1C), confirming that both INS-4 and INS-6 proteins, still expressed in these animals, are not functional in the absence of CAPS/UNC-31. The *daf-28(-)*;*unc-31(-)* dauers were persistent, in agreement with the known role of INS-6 in dauer exit^67^. Unexpectedly, *ins-4/6(-);unc-31(-)* animals developed normally (Fig. 1C), showing that there was sufficient DAF-28 function to prevent dauer entry in these animals, despite the absence of CAPS/UNC-31.

To verify that this was not due to some peculiar property of the *unc-31(e928)* null allele, we repeated the experiment with another, hypomorphic *unc-31(e169)* allele^87^. As with *unc-31(-)*, introduction of the *unc-31(hyp)* allele caused inappropriate dauer entry in *daf-28(-)* animals, but not in *ins-4/6(-)* animals (Sup. Fig. 1A).

To confirm that the lack of dauer entry in *ins-4/6(-);unc-31(-)* strain was indeed due to the ability of DAF-28 protein to signal normally in the absence of UNC-31, we used our functional DAF-28::mCherry transgene^76^. We found that the ability of DAF-28::mCherry to promote development and prevent dauer entry of *daf-28(-)*;*ins-4/6(-)* animals, lacking all three pro-growth IGFs, was as effective in the absence of UNC-31 (Fig. 1C) as it was in its presence (Fig. 1A).

To similarly confirm that the abnormal dauer entry in *daf-28(-)*;*unc-31(-)* animals was due to the inability of either INS-4 or INS-6 to function, we took advantage of the recently reported INS-6::mCherry transgenic protein, expressed from its cognate *ins-6* promoter^72^. INS-6::mCherry partially rescued the inappropriate dauer entry in the *ins-4/6(-);daf-28(-)* background (Fig. 1A), indicating that it is functional. The lower rescue is in agreement with DAF-28 having a more prominent role than INS-6 in the ASI for promoting development and preventing dauer, while INS-6 is stronger in signaling recovery from dauer, in the ASJ neuron^67,72^. Indeed, when scored a day later, *ins-4/6(-);daf-28(-)* animals expressing INS-6::mCherry exhibited increased dauer recovery (increase from ∼11% recovery in non-transgenic animals to ∼53% in INS-6::mCherry animals (Sup. Fig. 1B (Enter-exit)), in addition to the decreased initial activation of dauer (from ∼3% normal development in non-transgenic to ∼33% in INS-6::mCherry animals (Sup. Fig. 1B (Norm. dev.)). Having confirmed the functionality of INS-6::mCherry protein, we tested its dependence on UNC-31. We found that INS-6::mCherry expression failed to rescue development in the *ins-4/6(-);daf-28(-);unc-31(-)* genetic background, with only 1 of 489 animals avoiding the inappropriate dauer entry (Fig. 1C); these dauers retained their persistence.

Therefore, the ability of the three IGF-like growth factors, DAF-28, INS-4, and INS-6, to signal commitment to reproductive development in response to pro-growth environment is differentially dependent on CAPS/UNC-31. While INS-4 and INS-6, as expected, are UNC-31 dependent, DAF-28 functions normally in the absence of UNC-31, as confirmed both for the endogenous protein at genetic level, and for the fluorescent fusion protein. These data may also help explain the absence of dauer-constitutive phenotype in *unc-31* mutant animals under pro-growth conditions (through the continuing function of DAF-28), as well as the known defect in recovery of stress-induced *unc-31* dauers^87^ (through the failure of INS-6 function). Interestingly, it was previously observed that killing the ASI neuron, but not the ADF, ASG or ASJ neurons, does abnormally trigger dauer in *unc-31* mutants^87^; this is consistent with the ASI being the main source of UNC-31-independent signal for continued development.

### DAF-28, like INS-6, is routed into the regulated secretory pathway

One potential explanation of DAF-28’s independence of UNC-31 is that it does not utilize the regulated pathway in neurons, and instead is secreted *via* the default pathway. We used the functional DAF-28::mCherry protein to directly test this. Unlike the peptides and proteins that are constitutively released through the default pathway, neuronal secretory cargos in the regulated pathway are actively transported in vesicles by appropriate motor proteins to their release sites, where they are stored until their release is triggered by neuronal activity^58,88,89^. In *C. elegans*, most DCV peptides and proteins therefore exhibit a strongly polarized accumulation in presynaptic regions in axons, in a punctate pattern representing their release sites^4,79^, and, like in vertebrate neurons, require the anterograde axonal kinesin KIF1A/UNC-104 for this localization^1,56,90,91^. We first confirmed that DAF-28 selectively localized to axonal release sites. We have previously reported that DAF-28::mCherry localizes in a punctate pattern along the ASI axons^6^. In addition to the ASI, *daf-28* is expressed in a number of head and tail neurons^71^, and closer examination confirmed that DAF-28::mCherry accumulates in the axons of all of these neurons (Fig. 2B,D). We observed multiple puncta within synapse-rich axonal varicosities of head neurons (Fig. 2B’ and Sup. Fig 2A,B), indicating accumulation at putative release sites.

**Fig. 2.**
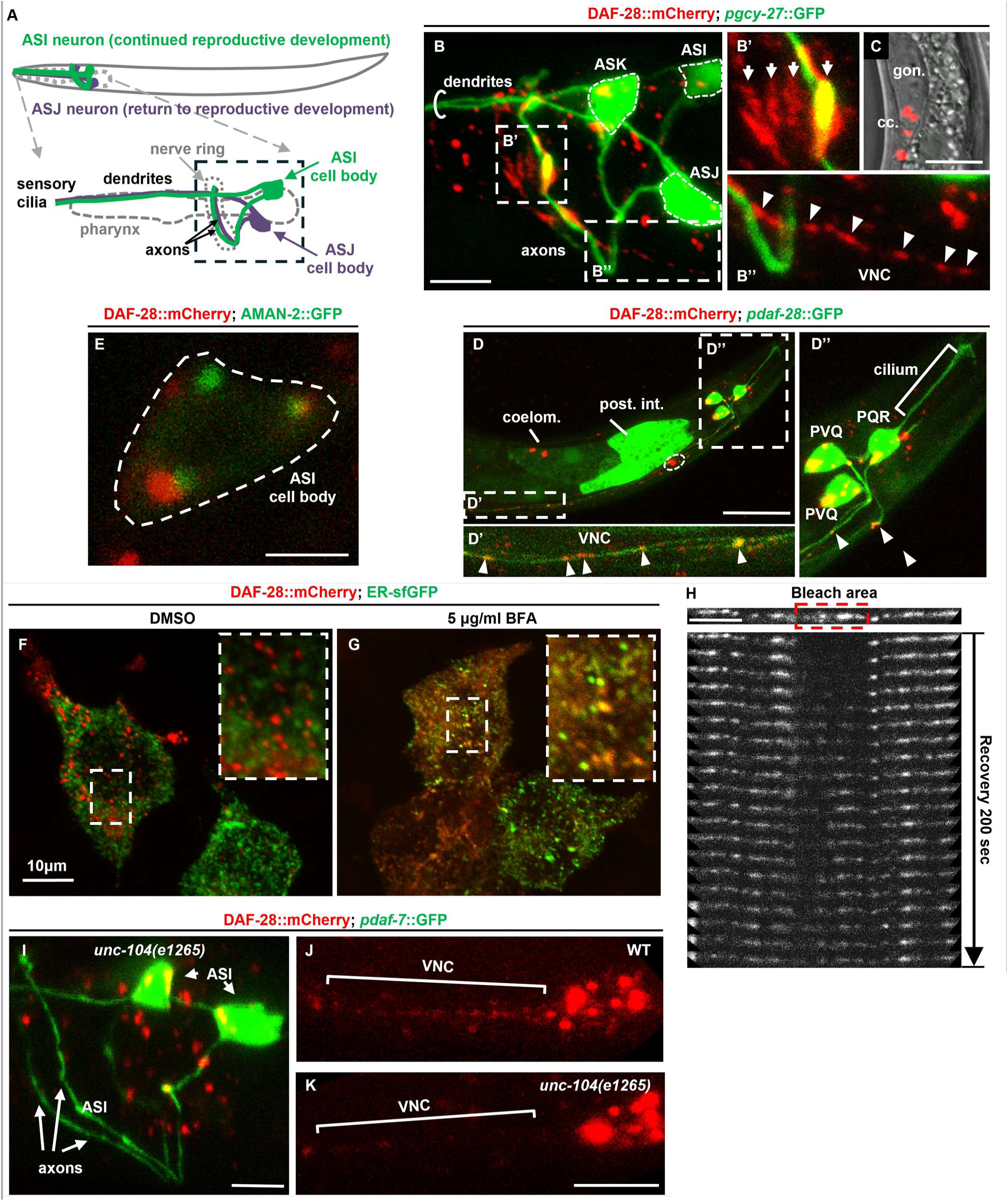
DAF-28 uses the regulated secretion pathway in neurons. **A.** Schematic of the ASI and ASJ neurons. Nerve ring is a bundle of axons of sensory and interneurons, rich in chemical and electrical synapses. Anterior is to the left, dorsal up. Punctate square outlines head area containing axons in the nerve ring and neuronal cell bodies. **B.** DAF-28::mCherry protein (red) is expressed in a number of neurons in L2 larva and localizes to the axons within the nerve ring. Green: GFP driven by *pgcy-27* promoter, used to visualize the ASI, ASJ, and ASK sensory neurons. Shown is the axonal area, as indicated in **A**; examples of entire neurons including the dendrites are in Sup. Fig. 2. **B’ and B”** are enlargements of the areas indicated in **B**, showing punctate accumulations of DAF-28::mCherry in varicosities of the nerve ring axons (arrows) and in the pre-synaptic ventral nerve cord (VNC) processes of tail neurons projecting to the nerve ring (arrowheads). **C.** mCherry signal in the coelomocytes (cc.) of the same animal. Shown is the second pair of coelomocytes, next to the gonad (gon.). **D.** DAF-28::mCherry localizes to puncta (arrowheads) along the VNC processes. *pdaf-28*::GFP-positive neurons are tentatively identified as PVQ and PQR; DAF-28 does not enter PQR cilia (**D’’**). **E.** Large DAF-28::mCherry puncta in the ASI cell body are adjacent to the the *cis*/medial-Golgi marker AMAN-2::GFP (green). **F,G.** *C. elegans* DAF-28::mCherry localizes to mobile vesicles that do not overlap with ER-sfGFP (green) in HEK293T cells (**F**). 6 hour treatment with 5 µg/mL BFA results in their strong co-localization (**G**). **H.** Localization of DAF-28::mCherry to the large axonal puncta (likely representing DAF-28 release sites) is dynamic. Area in the VNC, indicated by the red box, was photobleached, and the recovery of the fluorescent pattern was imaged with successive scans. **I-K.** DAF-28::mCherry does not localize to the nerve ring axons (**I**) or pre-synaptic VNC processes (**J and K**) in the absence of axonal kinesin KIF1A/UNC-104. **I** and **J** are the same animal. Scale bars for **B**, **H**, **I** and **K** = 5μm, for **D** = 20μm, and for **E** = 2μm.

This localization pattern in the head neurons was stereotyped, as illustrated by comparing left and right neurons in the same animal (Sup. Fig. 2B). Similarly, DAF-28::mCherry was found in punctate pattern in the axons of several tail neurons projecting into the nerve ring (Fig. 2B’’ and D’), tentatively identified as the PVQ interneuron and PQR neuron. DAF-28::mCherry was excluded from both the dendrites of the head neurons and the sensory cilium of the PQR neuron (Fig. D’’ and Sup. Fig. 2A). Thus, DAF-28::mCherry exhibits polarized localization to presynaptic regions in multiple neurons.

While characterizing the DAF-28::mCherry subcellular localization, we were surprised by the absence of the typical cell-body localized signal representing the protein still in the endoplasmic reticulum (ER). Instead, we typically detected 2-3 large mCherry-positive puncta within the cell bodies of the ASI or tail neurons. These cell-body puncta likely represent the trans-Golgi, based on their adjacent localization to the *cis*/medial-Golgi marker AMAN-2::GFP^92^ (Fig. 2E). To understand the lack of ER localization of this protein, we expressed DAF-28::mCherry in mammalian HEK293T cells. As in the worm neurons, we observed a punctate vesicular pattern that did not overlap with the co-expressed ER marker, ER-sfGFP (Fig. 2F). However, inhibiting ER-to-Golgi trafficking by 6hr treatment with Brefeldin A (BFA) resulted in clear co-localization of DAF-28::mCherry with ER-sfGFP (Fig. 2G). Since chromophore maturation in mCherry is relatively slow^93,94^, these data indicate that DAF-28::mCherry protein exits the ER before the chromophore in mCherry tag matures and becomes fluorescent. Indeed, DAF-28::mCherry in mammalian cells was secreted into the media, and secretion was disrupted by BFA treatment (Sup. Fig. 2C).

To confirm vesicular movement, we imaged the recovery of punctate fluorescence after photobleaching. We chose to image axons of the tail neurons, due to their planar geometry in the ventral nerve cord (VNC). The large DAF-28::mCherry puncta indeed recovered within minutes after photobleaching (Fig 2H); we also detected the movement of fluorescent vesicles (Sup. Fig. 2D and Sup. Movie 1). Finally, we confirmed that DAF-28 localization to axonal/pre-synaptic compartments depends on KIF1A homologue UNC-104 in both the head and the tail neurons (Fig. 2I-K and ref^6^), using the loss-of-function allele *unc-104(e1265)*^95^. We found no detectable DAF-28::mCherry signal in the axons in the nerve ring, and a loss of signal in the VNC, in *unc-104(e1265)* animals. Together, these data show that DAF-28 accumulates at pre-synaptic/axonal release sites, and its localization to these sites requires active vesicle trafficking by the KIF1A/UNC-104 motor.

Although the CAPS-dependent function of INS-6 is already consistent with its localization *via* the regulated pathway, we wanted to confirm this assumption. *ins-6* is transiently induced in the ASI in well-fed L1/L2 larvae, and is strongly induced in the ASJ in animals recovering from starvation or from dauer arrest^67,68,71,72^.

Surprisingly, we were unable to visualize INS-6::mCherry protein in the axons of the ASI neurons of well-fed larvae, despite detecting the co-injected transcriptional reporter (Sup. Fig. 3A,C). As we show later, this was due the efficient release of INS-6 from the axons, taking its fluorescence below the detection limit. However, we did detect strong accumulation of INS-6::mCherry within the axons of the ASJ neurons, in animals recovering from dauer (Sup. Fig. 3B), and in ASI axons of late L1/early L2 larvae recovering from starvation (Sup. Fig 3E). This axonal localization was lost in the presence of *unc-104(e1265)* allele (Sup. Fig. 3E,F).

**Fig. 3.**
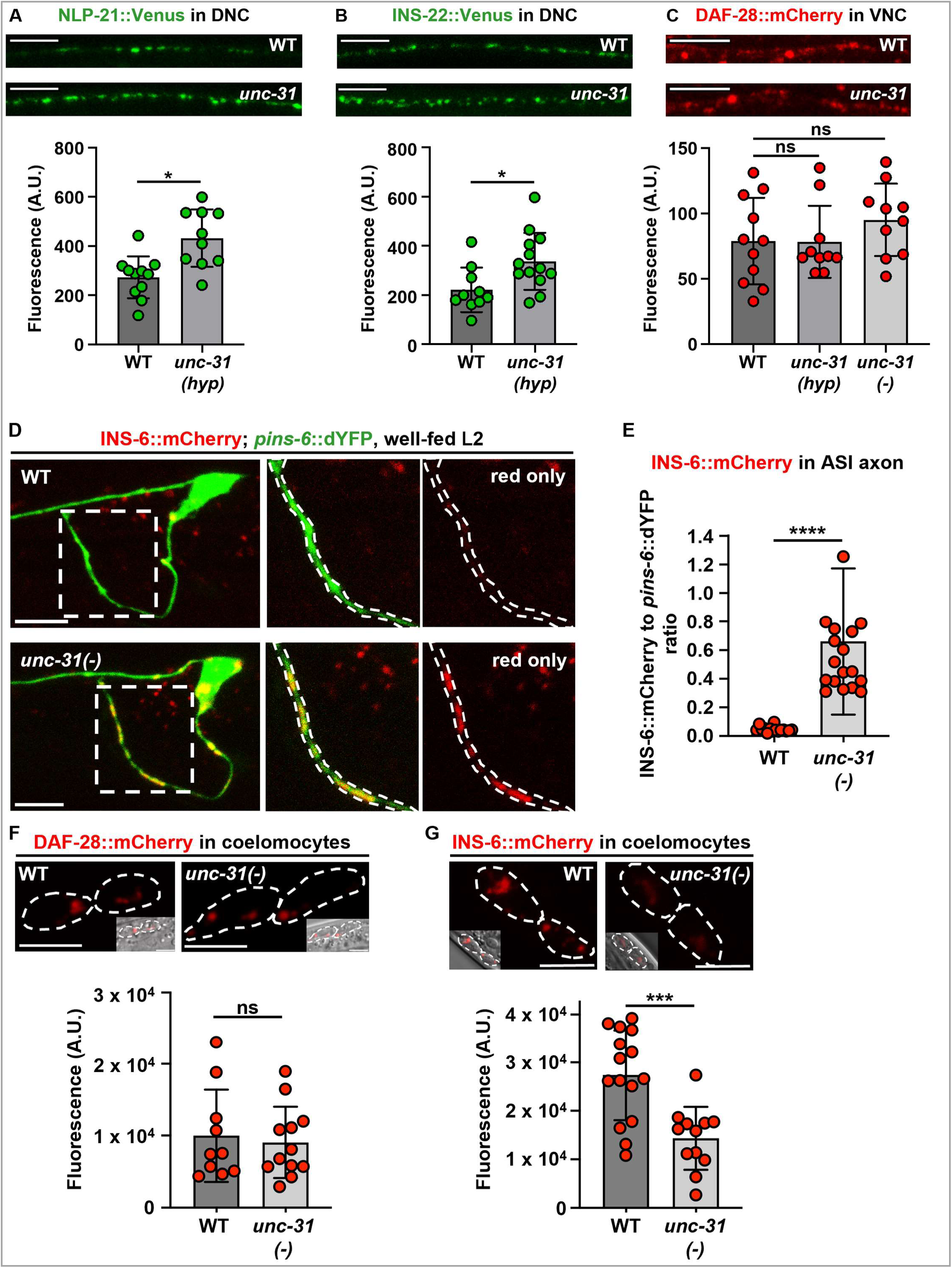
DAF-28 does not require CAPS/UNC-31 for axonal release, unlike model DCV cargos or INS-6. **A,B.** NLP-21::Venus and INS-22-Venus in motor neurons accumulate in axons projecting to the dorsal nerve cord (DNC), in a punctate pattern. *unc-31(hyp)* mutation*(e169* allele) results in significantly increased axonal accumulation of both NLP-21 and INS-22. **Top:** Confocal projections of the DNC; **bottom:** quantification of axonal GFP fluorescence. NLP-21::Venus: wild-type n=11, *unc-31(hyp)* n=10 animals; INS-22::Venus: wild-type n=10, *unc-31(hyp)* n=13 animals. **C.** DAF-28::mCherry accumulation (red) is similarly imaged (**top**) and quantified (**bottom**) in axons of tail neurons, in the ventral nerve cord (VNC). Mutation (*hyp, e169* allele) or loss (*e268* allele) of UNC-31 did not affect DAF-28::mCherry’s axonal accumulation.

Approximately half of the wild type animals (14/28) exhibited axonal localization compared to none (0/20) of the *unc-104(-)* animals (*p* = <0.0001, Fisher’s exact test (two-sided) C.I. 95%). At the same time, the ASI cell bodies in *unc-104(-)* animals contained enlarged puncta (Sup. Fig. 3D,F, arrows), suggesting that, in the absence of the UNC-104 motor, INS-6::mCherry accumulated in enlarged vesicles in the cell body. Therefore, both DAF-28 and INS-6 require KIF1A/UNC-104 for their localization to the presynaptic/axonal release sites.

### CAPS/UNC-31-independent function of DAF-28 is explained by its axonal release properties

Because *daf-28* is expressed in many neuronal and non-neuronal cells^96^, our genetic data (Fig. 1) do not unambiguously show that the difference in the ability of DAF-28 and INS-6 proteins to function in the absence of UNC-31 is due to their differential release by neurons. Furthermore, the differential UNC-31-dependence could be due to a role of UNC-31 in a process other than vesicle docking and release, as has been suggested^97,98^. To test this, we took advantage of the previously developed assay in worm neurons, that measures levels of fluorescently-labelled vesicle cargo in axons upon depletion of a release factor; increased fluorescence at the axonal release sites, and the corresponding decreased uptake into the scavenging cells (coelomocytes), indicates decreased release of the cargo^4,57,84^. As expected, the known DCV cargoes NLP-21::Venus and INS-22::Venus, expressed in a subset of motor neurons^4,79^, significantly accumulated in the dorsal nerve cord (DNC)-projecting axons in *unc-31(hyp)* animals (*e169* allele): the mean axonal fluorescence of NLP-21::Venus and INS-22::Venus increased by ∼1.6 and ∼1.5 fold (Fig. 3A,B), respectively, consistent with published data^4,99^. In contrast, DAF-28::mCherry did not measurably accumulate in the nerve ring–projecting axons of tail neurons in either *unc-31(hyp)* (*p*=0.97, two-tailed *t*-test) or *unc-31(-)* (*p=*0.24) animals (Fig. 3C).

Tail neurons were chosen for the quantitative assessment because their axons are planar, similar to the motor neurons, and because the curved axons of the multiple head neurons that express *daf-28* bundle together. For an additional, independent confirmation that DAF-28::mCherry is released normally in the absence of UNC-31, we measured its accumulation in coelomocytes (Fig. 3F). mCherry fluorescence levels did not change in *unc-31(-)* animals relative to the wild type (*p*=0.9229). These data are consistent with CAPS/UNC-31-independent release of the DAF-28 protein from its axonal release sites. Finally, because another Ca^2+^-dependent protein MUNC-13/UNC-13, which is normally required for synaptic vesicle release^100,101^, has been suggested to contribute to the DCV release^102,103^, we tested if it was needed for the axonal release of DAF-28. We found that neither NLP-21 nor DAF-28 accumulated in the axons of *unc-13(-)* animals (Sup. Fig. 4).

**Fig. 4.**
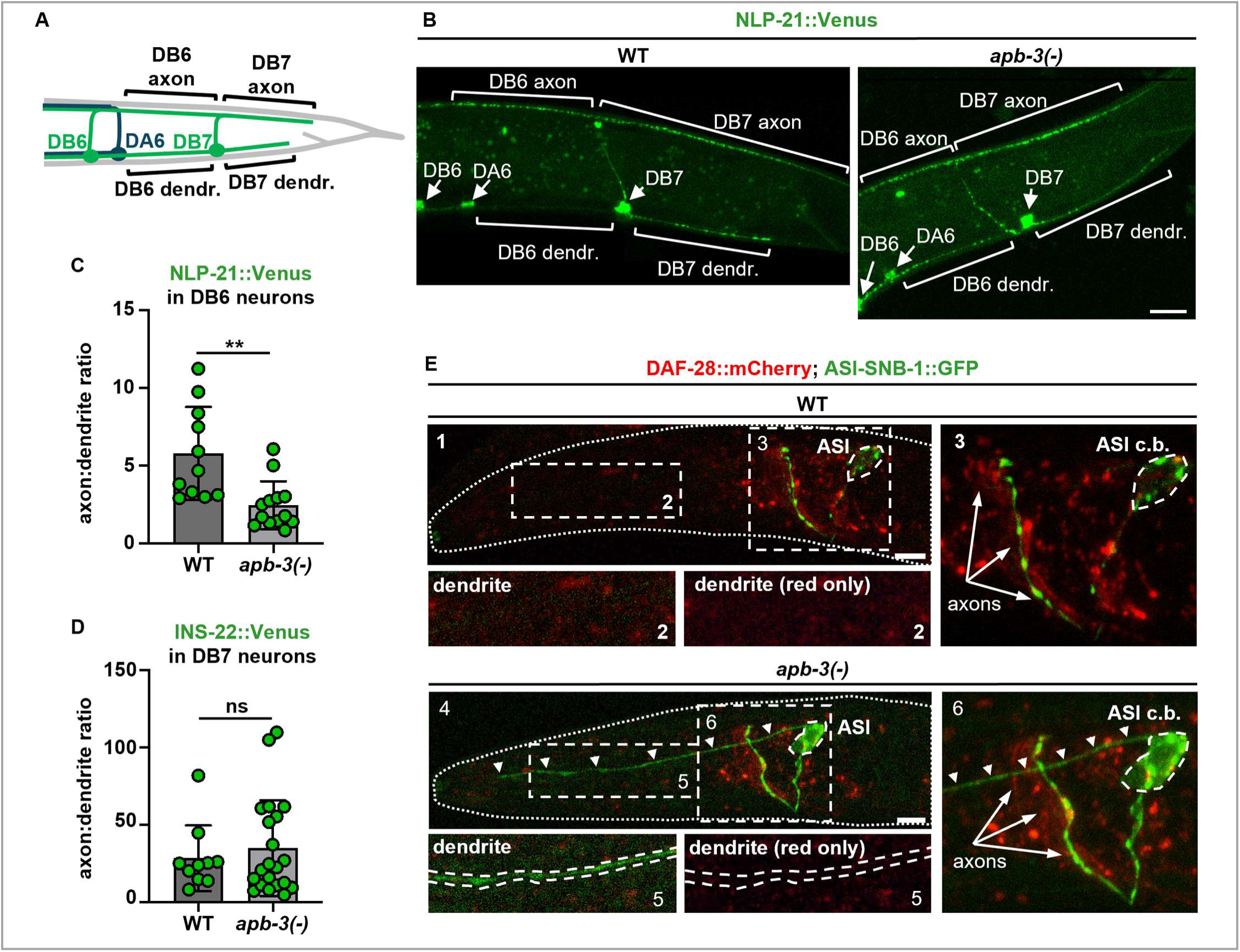
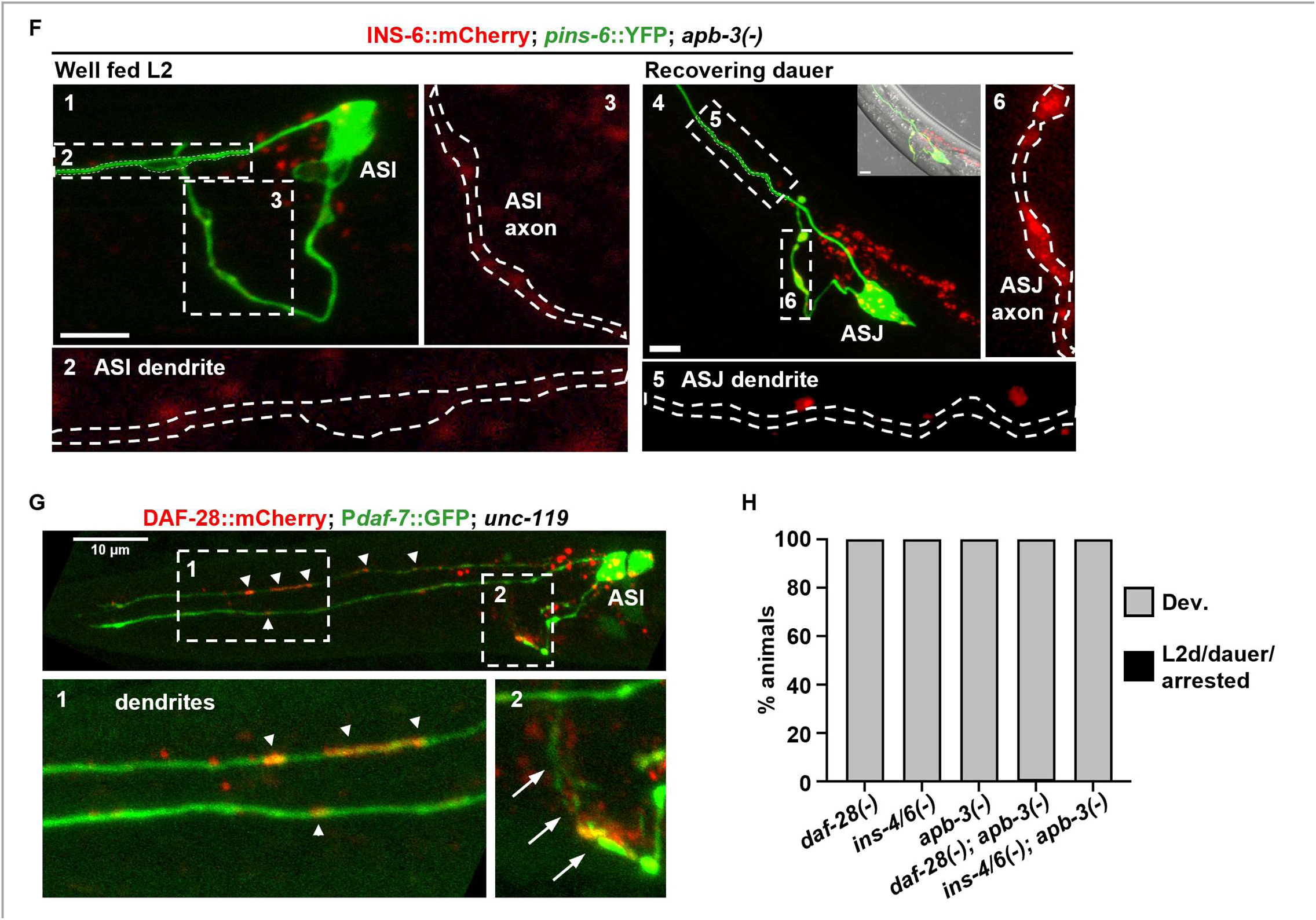
Axonal localization of insulin/IGF-like growth factors is independent the clathrin adaptor AP-3/APB-3, while that of neuropeptide NLP-21 is APB-3-dependent. A. Schematic of DA6 (blue), DB6 and DB7 (green) motor neurons, expressing NLP-21::Venus or INS-22::Venus. B. Confocal projections of NLP-21::Venus in DB6 and DB7 neurons; cell bodies indicated with arrows. C. Quantitation of the ratio of axonal to dendritic fluorescence of NLP-21::Venus in DB6 neurons, in wild-type (n=11) and *apb-3(-) (ok429)*(n=13) animals. The axonal preference seen in WT animals is strongly decreased in the absence of APB-3. DB7 neuron is in Sup. Fig. 5A. D. Quantitation of the ratio of axonal to dendritic fluorescence of INS-22::Venus in DB7 neurons of wild type (n=10) and *apb-3(-)* (n=21) animals. Loss of APB-3 did not affect the preferential axonal localization. Images are in Sup. Fig. 5B. E. DAF-28::mCherry (red) and ASI-specific SNB-1::GFP (green) in the wild-type and *apb-3(-)* animals. SNB-1::GFP is restricted to the synaptic region of the ASI axon in the wild type, but is non-polarized and readily detectable in dendrites (arrowheads) in *apb-3(-)* animals; DAF-28::mCherry remails exclusively axonal in the same *apb-3(-)* animal. Panels 2 and 5 show close-up of indicated dendritic area, with brightness increased 2-fold relative to the axonal areas (3, 6). F. INS-6::mCherry (red) remains excluded from dendrites in *apb-3(-)* animals in both ASI neurons (well-fed L2) and ASJ neurons (recovering dauer). dYFP expressed by the *pins-6*::dYFP array (green) is used to outline axons (panels 3 and 6) and dendrites (panels 2 and 5). G. DAF-28::mCherry (red) and ASI-specific GFP (*pdaf-7*::GFP, green) in an *unc-119* background. Arrowheads: DAF-28::mCherry mislocalized to dendrites; arrows: axonal DAF-28::mCherry. H. Endogenous DAF-28, INS-4, and INS-6 are functional in *apb-3(-)* animals*; s*cored as in Fig. 1. Significance in C, D: *t*-test with Welch’s correction, α=0.05. Scale bars in E and F = 5µm, in B = 10µm.

INS-6::mCherry protein is only expressed in either the ASI or ASJ neurons^72^; although their axons are non-planar, expression in only one neuron allowed measurements in the 3D curved axons. We used the dYFP signal of the short-lived *ins-6* transcriptional reporter (p*ins-6*::dYFP), included in the same transgene^72^, to produce a mask in which to measure mCherry fluorescence (Fig. 3D, punctate outlines), and normalized mCherry signal to that of dYFP within the mask. dYFP was similar in the wild type and *unc-31(-)* animals, suggesting no expression changes (see data supplement). While INS-6::mCherry fluorescence was nearly undetectable in the ASI axons of well-fed L2 larvae (Fig. 3D, top panel), as mentioned above (Sup. Fig 3A,C), we observed a remarkably strong accumulation (∼14 fold, *p*<0.0001, two-tailed *t-*test) of punctate INS-6::mCherry fluorescence in the ASI axons of well-fed *unc-31(-)* animals (Fig. 3D, bottom panel, and E). This large accumulation points to INS-6 being efficiently released in wild type animals, explaining the low fluorescence in their axons; the efficient release in the wild type is also supported by the presence of INS-6::mCherry signal in the coelomocytes and amphid sheath glia (Fig. 3G and Sup. Fig. 3A,B), which take up excess of released proteins. Finally, unlike with DAF-28::mCherry, we detected a significant (∼2 fold, *p=*0.0004, two-tailed *t-*test) decrease in INS-6::mCherry in the coelomocytes of *unc-31(-)* versus wild type animals (Fig. 3G).

Wild-type n=11, *unc-31(hyp)* n=10; *unc-31(-)* n=10 animals. **D.** Loss of UNC-31 resulted in dramatic increase of INS-6::mCherry in the ASI axons of well-fed L2 larvae. **Left panels:** confocal projection of the axonal area (as in Fig. 2A) of representative left ASI neuron; **right panels:** close up on the synaptic region of the axon. **E.** Ratio of INS-6::mCherry to dYFP fluorescence within the synaptic regions of ASI neurons in well-fed L2 larvae. As dYFP is co-induced with the INS-6::mCherry, the ratio is independent of changes in expression. Wild-type n=19, *unc-31(-)* n=18. **F,G.** Uptake of DAF-28::mCherry and INS-6::mCherry as indirect measure of their release from neurons. Loss of UNC-31 results in decreased coelomocyte uptake of INS-6::mCherry **(G)**, but not of DAF-28::mCherry **(F)**. DAF-28::mCherry: wild-type n=10, *unc-31(-)* n=12; INS-6::mCherry: wild-type n=15 and *unc-31(-)* n=12. All data were analyzed by unpaired *t*-test with Welch’s correction, α=0.05. Scale bars for **A-C** = 10µm, scale bars for **D,F,G** = 5µm.

Therefore, two independent assays show that the axonal release of INS-6, but not of DAF-28, depends on UNC-31 in the ASI neuron, suggesting that the ASI neuron is able to independently control the release of two structurally and functionally related signaling molecules in response to sensing pro-growth environment.

### The clathrin adaptor protein AP3β/APB-3 indicates an additional point of divergence in the vesicle pathways

One possible way by which peptides or proteins can be differentially released from the same neuron is if they are contained in non-overlapping populations of vesicles, for example for localization to distinct release sites. How cargos are selected into vesicles destined for different locations within an axon is not known, but the mechanisms for cargo selection into axonal *vs* dendritic vesicles have been elucidated, at least for membrane proteins^53^. These mechanisms rely on interaction of specific sequence motifs in the cytosolic tails of membrane cargo proteins with clathrin adaptors; sequences driving preferential interaction with AP3 rather than AP1 adaptor complexes result in enrichment of the corresponding cargos into axonally rather than dendritically targeted vesicles^53,104,105^. The soluble (non-membrane) cargoes are, in contrast, contained within vesicle lumen, and do not have direct access to the cytosolic machinery; their vesicle selection mechanisms are still undefined. However, it has been proposed that soluble cargoes may localize to the correct vesicles *via* cargo receptors - transmembrane proteins bridging between the vesicle’s lumen and the cytosol, or *via* binding to other membrane proteins within the vesicle^49,51,52,106–108^. Therefore, we asked whether the neuropeptides and growth factors used in this study depend on AP3 for their localization to the axonal release sites.

In *C. elegans*, loss of β subunit (*C. elegans* APB-3) of AP3 clathrin adaptor complex causes multiple pre-synaptic membrane proteins, such as serotonin and glutamate receptor subunits SER-1 and GLT-4, to lose their polarized axonal localization, and instead to localize to both axons and dendrites^53^. Thus, we used the known *apb-3* null allele *ok429*^109^ (designated as *apb-3(-)*) to first test the known non-membrane DCV cargos, NLP-21::Venus and INS-22::Venus^4,79^. We used a previously established method to visualize polarization of fluorescent cargoes in the DB6 and DB7 motor neurons^110^, because their posterior projecting axons (in the DNC) and dendrites (in the VNC) are planar to each other, and also can be imaged without overlap from the neurites of other neurons (see scheme in Fig. 4A). We found that the strong axonal polarization of NLP-21::Venus protein in the DB6 neuron became compromised in *apb-3(-)* animals, with mislocalized fluorescent protein being detectable in the dendrites (Fig. 4B). The mean DB6 axon:dendrite ratio of 5.8 ± 3 (mean ± st.dev.) in wild type animals decreased to 2.4 ± 1.6 in *apb-3* mutants (*p*=0.002, two-tailed *t*-test) (Fig. 4C). These data show that AP3β/APB-3 clathrin adaptor protein is required for the polarized axonal targeting of NLP-21 neuropeptide in the worm motor neurons.

Surprisingly, NLP-21::Venus was already poorly polarized in the DB7 neuron in wild type animals (Fig. 4B and Sup. Fig 5A). The reasons for this are not clear at present, although we did notice consistently higher fluorescence in DB7 cell bodies, raising the possibility that the higher expression of this transgenic protein in DB7 saturates the axonal sorting machinery^111^.

In contrast to the neuropeptide, the normally strong polarization of INS-22::Venus in DB7 neurons was not affected by the loss of APB-3, and INS-22::Venus in *apb-3(-)* animals remained localized to the axons and essentially undetectable in dendrites (Fig. 4D and Sup. Fig. 5B). The mean DB7 axon:dendrite ratio was 28.3 ± 21.2 in wild type and 34.8 ± 31.1 in *apb-3(-)* animals (*p*=0.9834, two-tailed *t*-test). Thus, the axonal localization of INS-22 is AP-3-independent. Although INS-22::Venus fluorescence is relatively low, these data suggest that despite being released through the same CAPS-dependent mechanism as NLP-21, INS-22 may be targeted to the pre-synaptic axonal compartment of the same neuron in distinct vesicles.

We next tested whether DAF-28 and/or INS-6 axonal localization requires AP-3, using both fluorescent and functional assays. We introduced ASI-expressed synaptobrevin-1::GFP (SNB-1::GFP)^112^ into DAF-28::mCherry-expressing animals, to use as internal control for the loss of polarity, since this transmembrane synaptic vesicle protein has been recently reported to depend on APB-3 for axonal localization^113^. In the wild-type background, SNB-1::GFP localized to the pre-synaptic sites in ASI axons, and was completely excluded from the sensory dendrites (Fig. 4E, panels 1-3, green). Similarly, DAF-28::mCherry localized to the axons of multiple neurons and was excluded from the dendrites (Fig. 4E, panels 1-3, red), as before. SNB-1::GFP and DAF-28::mCherry partially overlapped in the ASI neuron, suggesting peri-synaptic localization of DAF-28. Loss of APB-3 in *apb-3(-)* animals caused SNB-1::GFP to lose its polarity and to inappropriately enter the dendrites of the ASI neurons; SNB-1::GFP also appeared more diffuse in the axon and filling the cell body in *apb-3(-)* neurons (Fig. 4E, panels 4-6, green). Strikingly, in the same animals, DAF-28::mCherry retained its strict axonal polarization in multiple neurons and did not enter the dendrites (Fig. 4E, panels 4-6, red). We further confirmed the normal axonal localization of DAF-28::mCherry and its absence from the dendrites in *apb-3* mutants, when we used the ASI-specific marker (*pdaf-7*::*GFP)* to visualize the entire ASI neurons (Sup. Fig. 5C). Of the individual ASI neurons scored in *apb-3(-)* mutants, 0/14 had DAF-28::mCherry mislocalization to the dendrites.

To make sure that we could detect DAF-28::mCherry mislocalization, if such existed, we introduced a genetic mutation known to generally disrupt neuronal polarity, *unc-119(ed3)*. UNC-119 is thought to function as a lipid chaperone and contribute to ciliogenesis and localization of myristoylated proteins, and was recently shown to maintain microtubules organization in neurons; loss of UNC-119 in worms disrupts axonal polarization of synaptic vesicles in PVD neuron^114–116^. We detected a striking loss of polarity of DAF-28::mCherry in *unc-119* mutant animals, with strong signal in both the dendrites and the axons (Fig. 4G), indicating that its lack of mislocalization in *apb-3* mutants was not due to a detection problem.

Finally, we tested AP-3-dependence of INS-6. Similar to INS-22::Venus and DAF-28::mCherry, we did not observe mistargeting of INS-6::mCherry to the dendrites in *apb-3* mutant animals (Fig. 4F). This was true both in the ASI neuron of well-fed L2 larvae, whereas we showed earlier, INS-6 is efficiently released and thus not detectable in the axons (Fig. 4F, panels 1-3), and in the ASJ neuron of recovering dauers, where INS-6 had strong signal in the axons and none in dendrites (Fig. 4F, panels 4-6).

To independently confirm the independence of DAF-28 and INS-6 of AP-3, and to test INS-4, we performed the functional test for endogenous proteins. We used the same assay as in Figure 1, and found that the loss of APB-3 did not impact the ability of either DAF-28 alone (in *ins-4/6(-);apb-3(-)* animals), or of INS-4 and INS-6 (in *daf-28(-);apb-3(-)* animals), to signal reproductive development (Fig. 4H). *apb-3(-)* animals did have variably penetrant phenotypes like shorter length (‘dumpy’ phenotype, *dpy*), dark intestine, motility defects, and embryonic lethality, possibly reflecting vesicle sorting defects in multiple tissues^117^, but these phenotypes were not changed by either *daf-28* deletion, or *ins-4/6* deletion. Thus, unlike the neuropeptide NLP-21::Venus and the transmembrane synaptic vesicle protein SNB-1::GFP, none of the four insulin/IGF-like growth factors examined in this study (INS-22, DAF-28, INS-4 and INS-6) required AP3β/APB-3 clathrin adaptor protein for their axonal polarization or function.

## Discussion

Our findings support the existence of distinct molecular pathways for the local release of signaling molecules from presynaptic/axonal compartment in *C. elegans* sensory neurons, and identify two points where these pathways differ in their molecular mechanisms. The most striking finding is that DAF-28, an IGF-like growth factor that is released by chemosensory neurons when they sense pro-growth environment, is released through a CAPS/UNC-31-independent mechanism, while its functional and structural homologues INS-4 and INS-6 are released through the canonical CAPS/UNC-31-dependent mechanism. The two mechanisms appear to operate at the same time and in the same neuron.

Previous data showed that loss of both CAPS1 and 2 proteins in mammalian neurons dramatically reduces DCVs release, while in *Drosophila* and *C. elegans* neurons, CAPS homologues dCAPS and UNC-31, respectively, are required for the release^4,57,83,84,118^. Thus, it has been generally accepted that CAPS dependence is a common characteristic of stimulus-dependent release of proteinaceous signaling factors, like neuropeptides and neurotrophins or growth factors. However, it is well established that DCVs can exist in the same neuron as distinct populations carrying different cargoes and different vesicle-associated proteins^29,30,32–34^, raising a question of differential regulation^35,36,119^. Our findings show that both CAPS-dependent and CAPS-independent pathways operate in the worm sensory neuron, and that cargo identity determines the pathway used for the axonal vesicle release.

The second molecular difference was identified in the mechanisms for targeting soluble (non-membrane) cargos to the axons. We found that axonal targeting of the neuropeptide NLP-21::Venus depends on AP-3 clathrin adaptor, similar to what is known for transmembrane cargoes^53,105^. In contrast, none of the four tested insulin/IGF-like proteins required AP-3 adaptor for their axonal localization, or for function.

The divergence of axonal vesicle localization and release mechanisms supports the idea that neurons that co-produce multiple protein- and peptide-based signaling factors can utilize layers of regulation to control their selective release. Selective vesicle release may in turn contribute to precision in neuronal responses to different sensory inputs, and to the ability of hard-wired circuits to enact different outcomes^36,37^. As these mechanisms are not well understood, model organisms that exhibit sophisticated responses to environmental stimuli, despite possessing small nervous systems, will offer a great opportunity to delineate their molecular composition.

### Exceptions to the CAPS/UNC-31 DCV pathway

Our data show that axonal release of DAF-28, INS-4, and INS-6 proteins is differentially dependent on the vesicle docking/release protein CAPS/UNC-31. We used two distinct approaches to determine UNC-31-dependence of these proteins – first, an established imaging assay that measures increased presence of the tested vesicle cargo at its release sites, with parallel decreased uptake into the scavenging cells, to indicate decreased release^4,57,84^; second, we developed a new functional assay (Fig. 1B). While both approaches showed that DAF-28 does not require UNC-31, the functional assay is particularly powerful, because it reflects the natural molecular and regulatory context, and obviates the limitations of using fluorescent fusion proteins, as discussed below. When continued development of young larvae was dependent only on the release of INS-1 or INS-6, due to the deletion of *daf-28*, loss of UNC-31 resulted >95% animals entering and remaining in dauer, despite the presence of abundant food and the presence of development-promoting signals (low population density and growth temperature). Because INS-4 and INS-6 act in the ASI, and possibly other neurons, for prevention of dauer entry, and in ASJ neurons to signal dauer exit^67,68,72^, this was the expected outcome if release of these two growth factors required UNC-31. In contrast, when continued development was dependent only on the release of DAF-28, due to the deletion of both *ins-4* and *ins-6* genes, loss of UNC-31 did not increase dauer or pre-dauer (L2d) formation, showing that DAF-28 continued to function normally^71^.

Other observations have indicated the presence of CAPS-independent pathway(s) in neurons. cAMP and its target PKA can increase release of DCV cargos, either by stimulating Ca^2+^ release from internal stores (primarily the endoplasmic reticulum) *via* IP3 receptor, often in the soma or dendrites^120^, or by phosphorylating multiple presynaptic exocytotic proteins^121–126^. The latter could arguably bypass the need for CAPS. Indeed, in *C. elegans* cultured neurons, a direct, Ca^2+^-independent cAMP/PKA action was able to overcome DCV docking defects caused by UNC-31 mutation, suggesting a PKA target that acts in parallel or downstream of UNC-31^127^. However, two *in vivo* studies in *C. elegans* motor neurons showed that UNC-31 was in fact required for activation of the Gα_s_ pathway^86^, and for cAMP-dependent release of NLP-21 neuropeptide^128^, placing cAMP/PKA upstream of UNC-31. cAMP/PKA thus may affect vesicle release *via* multiple mechanisms, and the potential CAPS/UNC-31-independent cAMP/PKA pathway will require careful further study.

Another possible alternative pathway may involve the sole worm homologue of Islet Antigen-2 (IA-2) and phogrin, IDA-1^129,130^. These proteins are transmembrane constituents of DCVs^131,132^, and although their exact molecular function is unclear, IA-2 and phogrin are implicated in regulating insulin secretion in β-cells^133,134^, and IA-2 and IDA-1 - in neuroendocrine secretion^135,136^. However, paradoxically, loss of IDA-1 suppressed, rather than enhanced, temperature-induced dauer formation in animals lacking UNC-31^137^, which is inconsistent with a positive role of IDA-1 in release of DAF-28 (or DAF-4 and DAF-6). If the putative function of IA-2 and phogrin in promoting secretion of insulins is conserved in the worm, it is possible that IDA-1 contributes to the release of pro-dauer signals such as INS-1 or INS-18.

DAF-28 appears to be different from other insulin/IGF-like proteins in its use of the alternative release pathway. In addition to INS-4 and INS-6 in this work, a more distantly related INS-22, and one of *C. elegans* insulin homologues, INS-1, were previously found to be UNC-31-dependent^59,85^. Although indirectly, this suggests that DAF-28 may be present in a different type of secretory vesicle than other insulin/IGF-like proteins. The reason for this is not immediately apparent, as all three IGFs examined here respond to the same sensory input (food), and signal the same organismal response (continued development), at least in the ASI neurons in young larvae. One possible reason for this divergence is that redundant release pathways may be used to ensure robust and precise decision to continue reproductive development *vs* enter dauer, similar to the use of parallel neuronal pathways^74^. On the other hand, these proteins do have other functions, and are also released from other neurons. For example, while INS-6 is released from the ASI to signal continued development in late L1/early L2 larvae, it’s main function is in the ASJ neuron, where it is released in dauer larvae in response to re-appearance of food, to signal dauer exit^67,68,71,72^. Since DAF-28, in contrast, has only a minor role in dauer exit^67^, the specific sensory signals and downstream consequences of their release may differ even within the narrow growth-*vs*-dauer signaling paradigm. Finally, these growth factors have other, quite variable, functions outside of dauer signaling. For example, INS-6 in the ASI neuron contributes to olfactory learning^138^, and in the ASJ neuron to the timing of oogenesis onset in adults^139^. It is possible that targeting these two proteins into distinct types of vesicles, and releasing them *via* distinct mechanisms, helps match organismal responses to specific but variable sensory inputs.

The uniqueness of DAF-28 in our assays is intriguing, because the mammalian IGF-1 growth factor was previously shown to use a divergent DCV pathway in murine olfactory bulb neurons^32,45^. While canonical DCVs associate with the major synaptotagmin isoform Syt1, vesicles containing IGF-1 were instead associated with Syt10; Syt10 then selectively controlled IGF-1 release^45^. Interestingly, IGF-1-positive vesicles were negative for a co-produced neuropeptide ANF and the DCV integral protein phogrin, while some components of the release machinery, such as complexins, were common^32^. The presence of different vesicle-associated proteins, such as synaptotagmins, may explain tuning the differential release of DCV cargoes to local variations in Ca^2+^ signals, for example depending on whether the local Ca^2+^ influx is stimulated by a voltage- *vs* ligand-gated channel, e.g. VGCCs *vs* TRPV1, as was seen in dorsal root ganglion neurons^140^, or originates from action potential–induced extracellular influx *vs* IP3-induced release from internal stores, as was shown for *Drosophila* insulin-like protein 2 in clock neurons^47^. Interestingly, olfactory social learning in mice, which requires Syt10-dependent IGF-1 release in olfactory bulb neurons, was associated with sustained Ca^2+^ transients^46^. The mechanisms defining which vesicles are paired with which vesicle-associated proteins are unclear, and to our knowledge, IGF-1 and Syt10 are the only selective pairing identified so far. It will be interesting in the future to test CAPS dependence of IGF-1-positive vesicles *vs* the canonical ones.

### Targeting neuropeptides and growth factors to axonal release sites

In addition to the differential release of the IGFs from the ASI neuron, we also found that the neuropeptide NLP-21::Venus is targeted to the presynaptic release sites differently from all four of the insulin/IGF-like growth factor proteins (ILPs) we tested. This was true in the same motor neuron for NLP-21 *vs* INS-22. The AP-3-independence of the four ILPs was confirmed in our study by both imaging and the functional assay. A larger sample of different neuropeptides and proteins will be needed to understand this difference. For example, because the fluorescent Venus tag is cleaved from the NLP-21::Venus precursor protein during neuropeptide processing, we do not know whether the divergence in AP-3-dependence is due to some potential differences in packing/aggregation within the dense cores for short peptides (NLP-21) *vs* larger proteins (ILPs). Indeed, neuropeptides and some proteins, like insulin in β cells, are known to use aggregation, often homotypic and/or dependent on high calcium and low pH (or zinc ions for insulin), or phase separation, for their entry into DCVs^52,141–143^. However, it is unclear whether all larger DCV cargoes, such as IGF-like proteins, can aggregate or undergo phase separation under these conditions: an *in vitro* NMR study found mature INS-6 remaining monomeric at pH as low as 3.6, even at 1.5 mM^144^. Furthermore, some of these larger cargos may not be amenable to reversible aggregation, and may even be harmed by these conditions^145^: the mammalian IGF-1 is prone to misfolding, and possess a thermodynamically stable non-native conformation^146,147^, and similarly, the mouse IGF-2 is strictly chaperone-dependent^148,149^.

Another possibility is that different soluble cargos interact with specific transmembrane proteins to localize selectively to different pre-synaptically targeted vesicles. If so, cytosolic interactions of these transmembrane proteins could explain the differential reliance of soluble cargos on AP3 adaptor complex^51,53^. Indeed, loss of AP-3 results in loss of polarity for membrane proteins (ref^53^ and Fig. 4E), and we observed the same for NLP-21, which would be consistent with dependence on AP-3-interacting transmembrane protein. The identity of such protein is still obscure, as only few cargo receptor-like interactions are known for the Golgi exit of soluble secretory cargos in neurons, including carboxypeptidase E for POMS, or Wntless for Wnt^49–52,150^. Finally, a separate question posed by our data is how do AP-3-independent cargos, like the four ILPs in our study, use the axonal kinesin and/or are targeted selectively to the axons. The nature and identity of interactors for axonal targeting of soluble cargos will thus need further study.

### Considerations in choosing a model for the neuronal localization and release studies

The differences in the selectivity of DCV targeting to the axon that we observed in DB6 *vs* DB7 motor neurons highlight the importance of a model used to study the localization and release mechanisms in neurons. NLP-21::Venus showed less-polarized axonal localization in DB7 than in DB6, while INS-22::Venus protein localized exclusively to the axon in both neurons. Although this could be due to neuron- or cargo-specific differences in the localization mechanisms, it is more likely that the transgenic proteins themselves contribute to this discrepancy. Recent data in mammalian neurons showed that even a subtle increase in expression levels of transgenically-expressed synaptic vesicle proteins disrupts their selective localization to axons, and results in some spillover into dendrites^111^. Because the fluorescence level of NLP-21::Venus in DB7 cell body is noticeably higher than in DB6, and higher than that of INS-22::Venus in both cells, it is likely that this level of expression already overwhelms the selectivity mechanisms.

Another important consideration for models of secreted peptides or proteins is selection of a fusion tag. We have previously found that unlike the exclusively axonally-localized mCherry-tagged DAF-28, a different available DAF-28 model - the DAF-28::GFP fusion protein accumulates in cell bodies and enters both axons and dendrites in ASI and ASJ neurons^76^. Testing for transgenic protein’s function resolved this discrepancy in favor of axonal localization: we found that DAF-28::GFP is not functional, and causes unconditional dauer entry when introduced into *daf-28(-)* background^76^, likely due to its oxidative misfolding^151,152^ and resulting ER stress in the ASI neuron^153^. DAF-28::mCherry, on the other hand, can efficiently prevent dauer entry even when all three pro-growth signaling molecules are deleted in *daf-28(-);ins-4/6(-)* animals (ref^76^ and Fig. 1A), arguing for a functional assessment of the tagged protein in specific neuron before its use for localization and release studies.

Because of these potentially confounding effects of the fluorescent fusions, we developed a functional assay for the corresponding endogenous proteins, taking advantage of the switch-like growth *vs* dauer phenotypic output, and the redundancy between the three IGFs in pro-growth signaling^65–68^. This was particularly important for confirming the CAPS/UNC-31 independence of DAF-28 release, since the lack of increase in axonal fluorescence, and of corresponding decrease in coelomocyte fluorescence, could be interpreted as loss of sensitivity due to transgene overexpression. In addition to the independent validation of the finding made with fluorescent fusion proteins, this assay provides a measure of the functional importance of the observed changes. For example, loss of UNC-31 resulted in ∼4.8 fold increase in mean fluorescence of INS-6::mCherry in the ASI axons, and a ∼1.9 fold decrease in coelomocytes; although both measures point to a contribution of UNC-31 to INS-6 release, they could not exclude a possibility that some INS-6 was still released in the absence of UNC-31, and that such still-released amount could still be sufficient for pro-growth signaling. However, the functional assay, by showing almost complete dauer entrance of *daf-28(-);unc-31(-)* animals, allows the conclusion that endogenous INS-6 (and INS-4) is unable to perform its pro-growth function in the absence of UNC-31.

## Materials and Methods

### Strains and genetics

Standard methods were used for worm culture and genetics^154^. Animals were kept at constant 20⁰C temperature throughout, unless specifically indicated. Animals were synchronized by picking gastrula-stage embryos. Plates for picking embryos were set with larval stage 4 (L4) or young adult animals, picked from populations kept in healthy uncrowded conditions for at least two generations; embryos were picked 1-2 days later.

The following strains were obtained from the *Caenorhabditis* Genetics Center (CGC): PS10640(*cmk-1*(*sy2277*[*cmk-1*::mKate2::AID*::3xFLAG)IV;*syIs875*[*ins-6p*::dYFP + *ins-6*::mCherry + *unc-122p*::GFP]), DA509(*unc-31(e928)*)IV, CB169(*unc-31(e169)*)IV, OH4348(*otEx2503*[*gcy-27(prom1)*::GFP + *rol-6(su1006)*]), HT2099(*(unc-119(ed3)*III;wwEx85[*daf-28p*::GFP + *unc-119(+)*]), FK181(*ksIs2*[p*daf-7*::GFP + *rol-6(su1006)*]), NM440(*unc-104(e1265)*II;*jsIs1*[pSB120(*snb-1*::GFP);pRF4 (*rol-6(su1006)*)]), OQ366(*tsp-6*(*syb4122*[*tsp-6*::wrmScarlet])X;*ulbEx78*[*F16F9.3p*::CFP]), KP3947(*nuIs183*[*unc-129p*::*nlp-21*::Venus + *myo-2p*::GFP]), KP3957(*nuIs190*[*unc-129p*::*ins-22*::Venus + *myo-2p*::NLS::GFP])X MT8004(*unc-13(n2813)*)I, RB662(*apb-3(ok429)*)I, CX3572(*kyIs105*[*str-3p*::*snb-1*::GFP + *lin-15(+)*]V;*lin-15B&lin-15A(n765)*X).

N2AM (wild-type) was a subclone of N2Bristol from Morimoto Lab. GJ2374 (*gjEx1295*[p*gpa-4*::*aman-2*::GFP + p*elt-2*::GFP) was a gift from Jansen Lab (Erasmus MC). Strain 2308(*daf-28(tm2308)*V) was from the National BioResource Project (Japan). ZM7963(*hpDf761*II;*daf-28(tm2308)*V) was a gift from Fang-Yen Lab (Ohio State University). The transgene *drxEx21*[p*daf-28*::*daf-28*::mCherry::*unc-54* 3’UTR] is described in ref^76^.

Crosses with *daf-28(tm2308)* were performed using fluorescent chromosomal markers^155^ from the following strains: EG8860(*unc-119(ed3)*III;*oxTi906*V) or EG7968(*unc-119(ed3)*III;*oxTi633*V). Crosses with *hpDf761*II deficiency (*aka. ins-4/6* deletion) were performed using *oxTi75*II marker isolated from EG8040(*oxTi302*I;*oxTi75*II;*oxTi411*III;*unc-119(ed3)*III;*him-8(e1489)*IV).

Where necessary, individual transgenes or mutations were isolated by crosses with the wild type (N2AM). For all strains, *unc-119(ed3)* mutation was removed from the strain background. All crosses were confirmed by PCR and restriction digest or sequencing, or by phenotype (if unambiguous).

### Developmental assays

Animals were grown on fresh plates, seeded with OP50 *E*. *coli*, at 20°C under non-crowded conditions for at least 2 generations prior to embryo picking, to avoid effects on dauer entry^65^. 20–40 young adult (YA) animals were allowed to lay embryos for 12-24 hours at 20°C. For rescue assays with non-integrated transgenes, only transgenic animals were used as parents; the non-transgenic progeny were used as sibling controls to the transgenic ones. For each genotype or condition, 100–200 gastrula-stage embryos were picked onto new plates and allowed to develop for 65–66 hours at 20°C. Resulting animals were then scored for their developmental stage: animals with embryos present *in uteri* were scored as reproductive adults; YA or late L4 stages (based on gonad development) as mildly delayed; and early L4 or earlier stages (mainly L2d and/or dauers) as severely delayed. L2d larvae were radially constricted to a lesser extent than dauers, had a uniformly dark intestine, and exhibited slow pharyngeal pumping^65^. All developmental assays were repeated at least three times; raw data are in the Supplemental Data Tables.

### Recovery from food deprivation

Animals were grown at 20°C on fresh plates seeded with OP50 *E*. *coli* until the food was completely consumed. Freshly food-deprived late L1/early L2 animals or dauer animals, as indicated, were picked onto fresh OP50 seeded plates and allowed to recover for 5-6 hours. The recovering L2 or dauer worms were used for imaging. Recovering animals were compared to well-fed L2 animals that were grown separately.

### Live imaging

Animals were mounted on 2% agar pads with 20mM sodium azide. Animals were imaged with Zeiss LSM700 confocal microscope at Cell Imaging Center, Drexel University, using 1.4NA 63x oil objective. Animals were kept in the paralytic no longer than 1.5-2 hours. For each transgene and cell type, lowest laser power producing specific signal was determined, to prevent phototoxicity and photobleaching, and used consistently across genotypes and conditions, unless specifically indicated. For quantitation, identical settings were used to collect images for comparisons across genotypes. 12-bit confocal stacks were reconstructed in Fiji(ImageJ)^156^ as 3D or maximum intensity projections, and where indicated, overlaid on single plane DIC images.

### Fluorescence quantification of neuropeptide and neuronal growth factor localization

Maximum intensity projections were used for quantification and were made in Fiji. Fluorescence was measured in a defined region of interest using ‘raw integrated density’ function; background for subtraction was measured from an identically sized region adjacent to the region of interest. At least ten animals were used per genetic background or condition.

#### Measuring fluorescence in the axons and dendrites of planar neurons

For axon:dendrite ratio, z-stacks through the dorsal and ventral nerve cords (DNC and VNC) in the posterior of the animals were used to generate maximum intensity projections. DB6 and DB7 neurons and their axons and dendrites were identified based on their cell body and commissure locations. Background correction was performed for the axons and dendrites separately. For tail neurons, an area of the VNC in the posterior of the animal was imaged.

To visualize vesicles, animals were rolled during mounting and those with VNC positioned near the coverslip were imaged. For fluorescence recovery after photobleaching (FRAP), an area of interest was selected as shown and bleached at full intensity with 555nm laser for 30-40 iterations. Timelapse images were collected in a single plane of 1.4 μm (FRAP) or 3.6 μm (Sup. Movie 1) thickness, centered on the VNC. For movies, AVI files were generated from timelapse images in Fiji and converted to MP4 files using open-source software (HandBrake).

#### Measuring coelomocyte fluorescence

The perimeters of the coelomocytes were traced using DIC channel; fluorescence within the traced region was then measured from the maximum intensity projection, using mCherry channel. Coelomocytes typically had higher fluorescence than neurons, laser power was adjusted to avoid signal saturation.

#### Measuring fluorescence in the synaptic regions of axons of the bilateral head neurons

The synaptic region of the ASI axon was outlined using the dYFP cell fill (the short-lived *ins-6* transcriptional reporter (p*ins-6*::dYFP), included in the same transgene as INS-6::mCherry^72^). Fluorescence intensity was measured in both the dYFP and mCherry channels in the outlined region and reported as a ratio. Only animals oriented in a lateral position were used and only the neuron closest to the cover slip was measured.

### Brefeldin A treatment

#### For imaging

HEK293T cells were transfected with pBUDCE4-[DAF-28(cDNA)::mCherry::V5+ER::sfGFP] plasmid using Lipofectamine 2000 (Invitrogen). 24hrs after transfection, the cells were treated with DMSO or 5μg/ml brefeldin A (Sigma) for 6hrs, fixed with 4% Paraformaldehyde (PFA), and imaged with the Zeiss LSM700 microscope at Cell Imaging Center, Drexel University, using 1.4NA 63x oil objective.

<u>For dot-bot: HEK293T cell, transfected as above</u>, were allowed to grow for 24 hours and treated with 2.5μg/ml Brefeldin A for another 24 hours. Cell supernatant was loaded into dot-blot apparatus (Bio-Rad) and transferred to nitrocellulose membrane (Bio-Rad). The membrane was probed with anti-V5 (Invitrogen) and IRDye-conjugated secondary antibody (Li-Cor), and scanned with Odyssey Infrared imager (Li-Cor).

### Statistics

All Chi-square, Fisher’s exact test, and unpaired *t*-test analyses were performed using Prism software (GraphPad, USA) and Microsoft Excel. For data with unequal variances, Welch’s correction was used. When means are shown, data are presented as mean+/-SD. α=0.05 was used for all analyses.

## Acknowledgements

We thank Drs. Gert Jansen and Christopher Fang-Yen for providing some *C. elegans* strains. Some strains were provided by the CGC, which is funded by NIH Office of Research Infrastructure Programs (P40 OD010440). Confocal experiments were conducted at Drexel University’s Cell Imaging Center, RRID:SCR_022689.

---

**Sup. Fig. 1.**
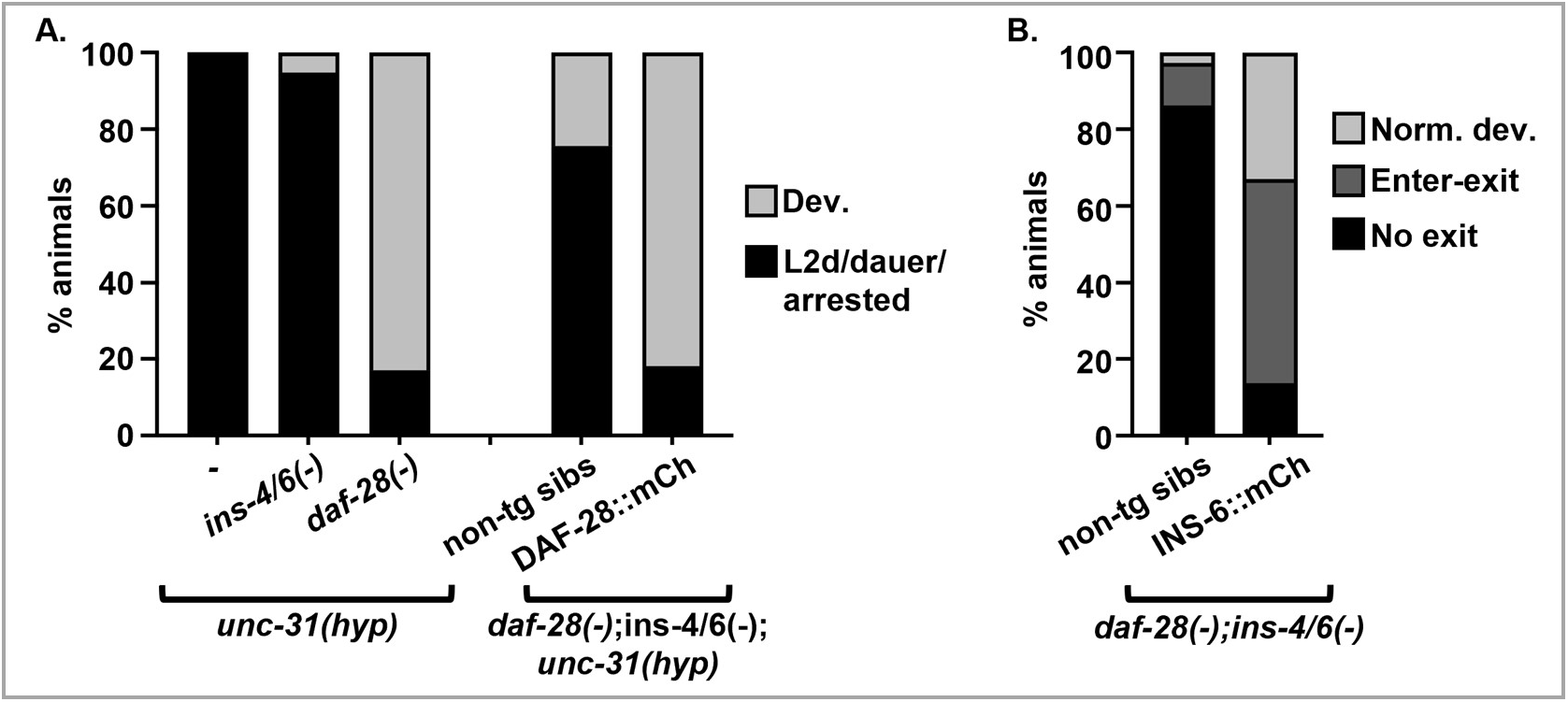
**A.** DAF-28 functional independence of UNC-31 is confirmed with an *unc-31(hyp)* allele *e169*. Scored as in Figure 1. **B.** INS-6::mCherry transgene is functional in signaling both reproductive development and dauer recovery. Development was scored ∼68-75 hours post-gastrula. **Norm. dev.** indicates animals that avoided dauer entry, **Enter-exit** – animals that recovered from L2d/dauer, **No exit** – animals still in L2d/dauer stage.

**Sup. Fig. 2.**
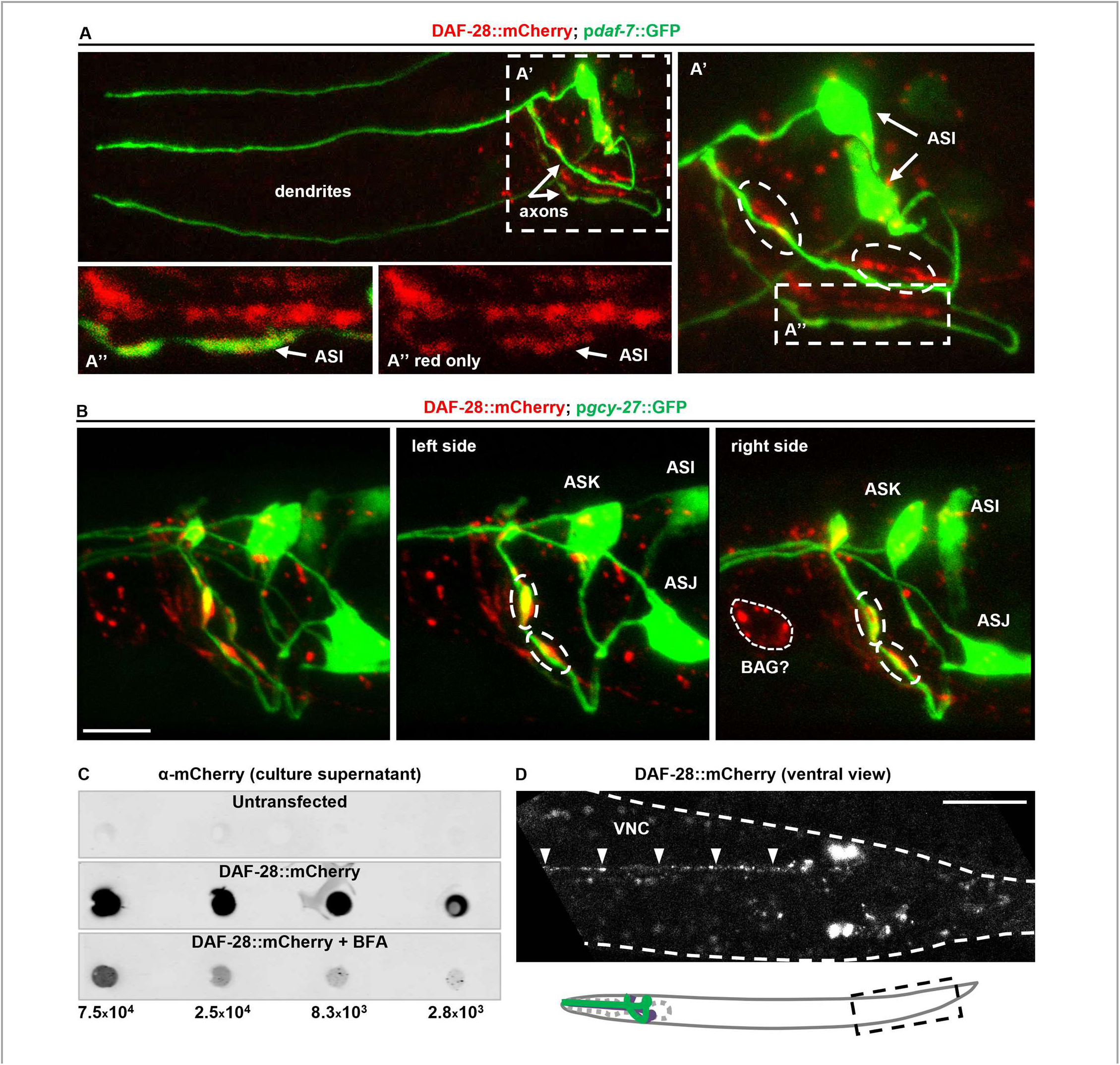
**A.** Confocal projection of the head of DAF-28::mCherry;*pdaf-7*::GFP animal, with both ASI neurons visible. DAF-28::mCherry protein (red) localizes exclusively to axons of the ASI and other neurons, and is excluded from dendrites. Green: GFP expressed from a *pdaf-7*::GFP;*rol-6* array, used to visualize the ASI. The third dendrite in A is from another animal. **A’** shows the axonal area. **B.** 3D confocal projections of axonal area of the animal in **Figure 2B,C**, showing both left and right neurons together (**left panel**) and separately; DAF-28::mCherry stereotypically accumulates in synapse-rich areas (ovals in A’ and B). Cell bodies of the ASI, ASJ, ASK, and likely Bag neurons are indicated. **C.** DAF-28::mCherry is secreted from mammalian cells. Dot blot of DAF-28::mCherry protein in media supernatant from transiently transfected HEK293T cells. The presence of DAF-28::mCherry protein in the media decreases dramatically following 24 hour incubation with 2.5 µg/mL BFA, indicating active vesicle-mediated release. Cell-number equivalents for the amount of media applied in each well are indicated at the bottom. **D.** Video of a section of the VNC (arrowheads) in the tail area, imaged from the ventral side. DAF-28::mCherry moves in bright puncta. A corresponding video file is **Sup. Video 1**. Scale bar for **B** = 5μm, for **D** = 20μm.

**Sup. Fig. 3.**
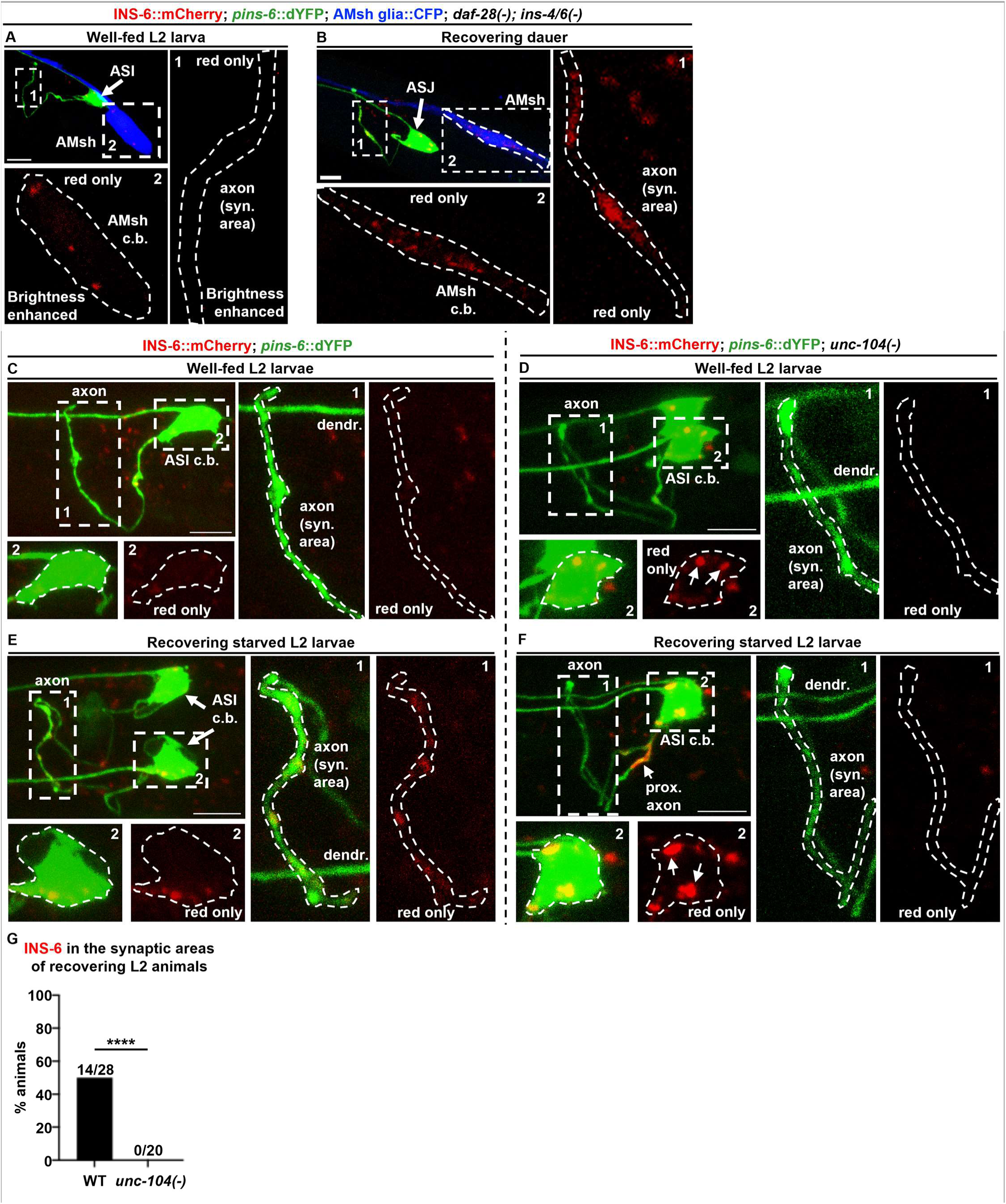
**A.** INS-6::mCherry (red) is undetectable in the axons of the ASI neuron in well fed L2 *daf-28(-);ins-4/6(-)* larvae, despite its induction (green). Its presence in the amphid sheath glia (AmSH, blue), due to uptake, suggests that it is released from the ASI neuron. *pins-6*::dYFP is present on the same array and drives expression of destabilized YFP (dYFP) from *ins-6* promoter; it indicates *ins-6* induction and allows visualization of the expressing neuron. **B.** INS-6::mCherry does accumulates in the axons of the ASJ neuron upon its strong induction in recovering dauers. This is consistent with a stronger signal in the AmSH glia, indicating higher release. **C,E.** In the wild type background, INS-6::mCherry is detectable in the synaptic regions of the ASI axons in L2 larvae recovering from starvation, but not in well-fed L2 (as in **A**). **D,F.** Loss-of-function of KIF1A/UNC-104 (*e1265* allele) results in the loss of axonal localization of INS-6::mCherry in recovering starved L2s (compare panels **E1** and **F1).** Instead, mCherry accumulates in enlarged puncta in cell bodies of both well-fed and recovering starved L2s, and in the proximal axon in the latter. **G.** Quantitation of the axonal localization defect penetrance in *unc-104(-)* animals. WT n=28, *unc-104(-)* n=20; Fisher’s exact test, two-sided, 95% confidence interval. Scale bars for **A-F** = 5µm.

**Sup. Fig. 4.**
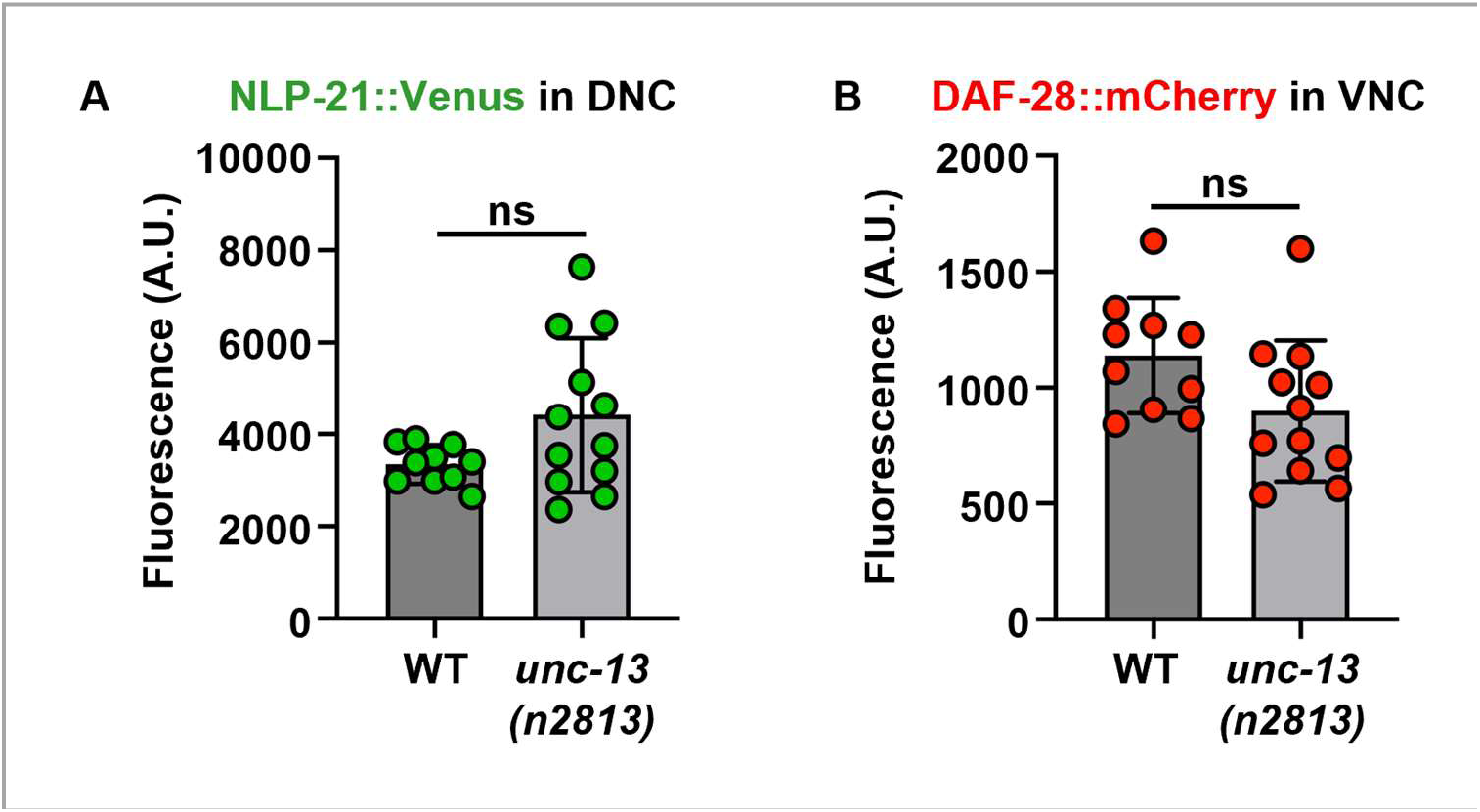
Axonal release of NLP-21 and DAF-28 is UNC-13-independent. A. Quantification of NLP-21::Venus protein in axons (in DNC) of wild-type (n=10) and *unc-13(n2813)* (n=12) animals. B. Quantification of DAF-28::mCherry protein in axons (in VNC) of wild-type (n=10) and *unc-13(n2813)* (n=12) animals. *t*-test with Welch’s correction, α=0.05.

**Sup. Fig. 5.**
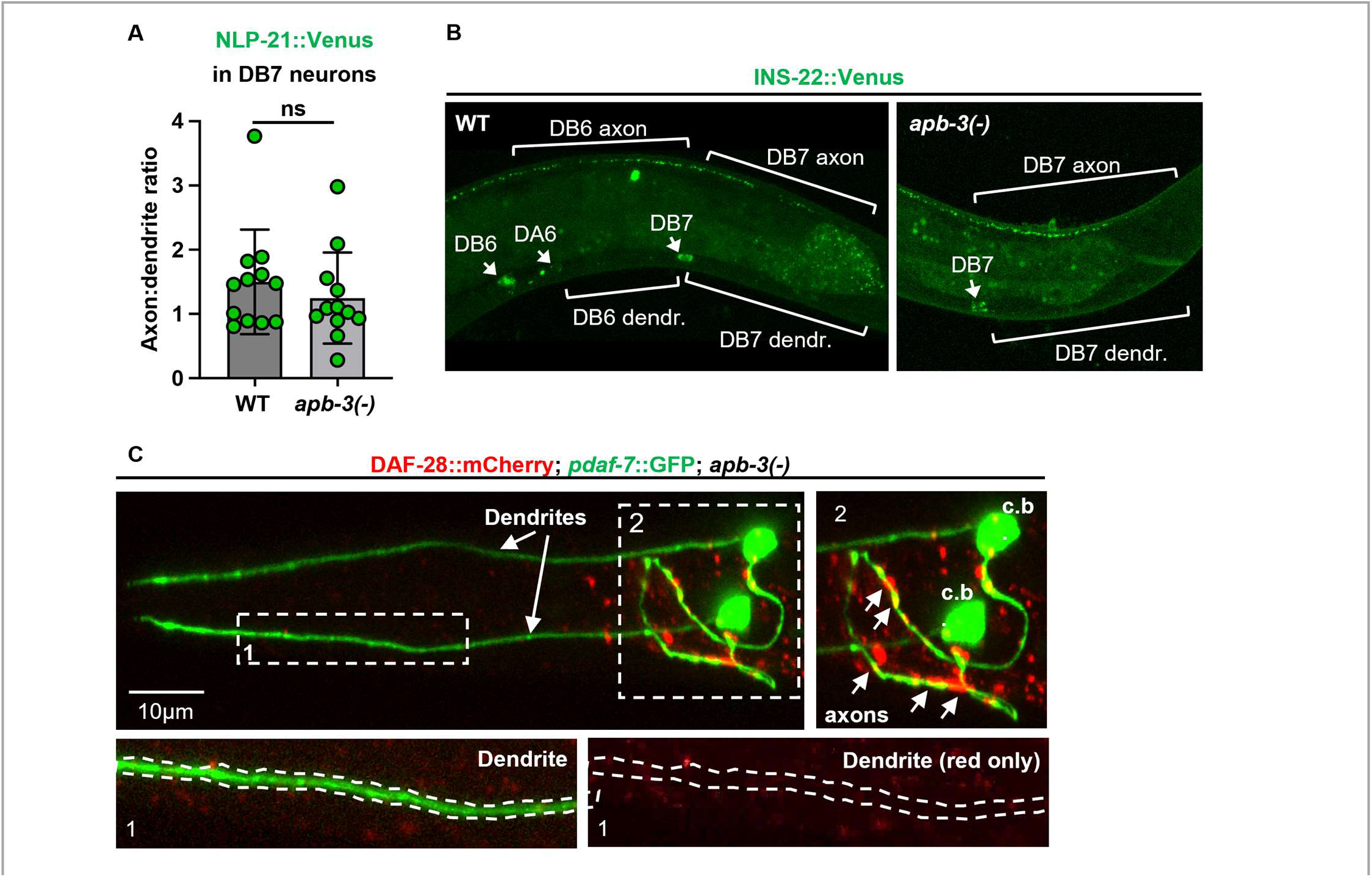
Insulin/IGF-like growth factors localize independently of APB-3. **A.** Quantification of NLP-21::Venus in the DB7 axons *vs* dendrites of wild-type (n=12) and *apb-3(-)* (n=12). **B.** Confocal projections of INS-22::Venus in DB6 and DB7 neurons; cell bodies indicated by arrows. **C.** DAF-28::mCherry (red) remains axonally localized and excluded from dendrites in an *apb-3(-)* background (*ok429* allele). Green: ASI-specific GFP (*pdaf-7*::GFP). Panel 1: dendrite area, with brightness increased 2-fold relative to the axonal area (panel 2); c.b.: ASI cell bodies; short arrows: DAF-28 accumulations in axons.

